# Ratio-based quantitative multiomics profiling using universal reference materials empowers data integration

**DOI:** 10.1101/2022.10.24.513612

**Authors:** Yuanting Zheng, Yaqing Liu, Jingcheng Yang, Lianhua Dong, Rui Zhang, Sha Tian, Ying Yu, Luyao Ren, Wanwan Hou, Feng Zhu, Yuanbang Mai, Jinxiong Han, Lijun Zhang, Hui Jiang, Ling Lin, Jingwei Lou, Ruiqiang Li, Jingchao Lin, Huafen Liu, Ziqing Kong, Depeng Wang, Fangping Dai, Ding Bao, Zehui Cao, Qiaochu Chen, Qingwang Chen, Xingdong Chen, Yuechen Gao, He Jiang, Bin Li, Bingying Li, Jingjing Li, Ruimei Liu, Tao Qing, Erfei Shang, Jun Shang, Shanyue Sun, Haiyan Wang, Xiaolin Wang, Naixin Zhang, Peipei Zhang, Ruolan Zhang, Sibo Zhu, Andreas Scherer, Jiucun Wang, Jing Wang, Joshua Xu, Huixiao Hong, Wenming Xiao, Xiaozhen Liang, Li Jin, The Quartet Project Team, Weida Tong, Chen Ding, Jinming Li, Xiang Fang, Leming Shi

## Abstract

Multiomics profiling is a powerful tool to characterize the same samples with complementary features orchestrating the genome, epigenome, transcriptome, proteome, and metabolome. However, the lack of ground truth hampers the objective assessment of and subsequent choice from a plethora of measurement and computational methods aiming to integrate diverse and often enigmatically incomparable omics datasets. Here we establish and characterize the first suites of publicly available multiomics reference materials of matched DNA, RNA, proteins, and metabolites derived from immortalized cell lines from a family quartet of parents and monozygotic twin daughters, providing built-in truth defined by family relationship and the central dogma. We demonstrate that the “ratio”-based omics profiling data, *i.e*., by scaling the absolute feature values of a study sample relative to those of a concurrently measured universal reference sample, were inherently much more reproducible and comparable across batches, labs, platforms, and omics types, thus empower the horizontal (within-omics) and vertical (cross-omics) data integration in multiomics studies. Our study identifies “absolute” feature quantitation as the root cause of irreproducibility in multiomics measurement and data integration, and urges a paradigm shift from “absolute” to “ratio"-based multiomics profiling with universal reference materials.

---

Multiomics profiling is a new approach where biomolecules across multiple omics layers including genomics, epigenomics, transcriptomics, proteomics, and metabolomics are fully measured, analyzed, and integrated from the same set of samples on a genome scale in terms of the number of measured features^1–3^. Multiomics profiling quantifies biologically different signals across complementary omics layers, therefore promises to demonstrate significant advantages over any single omics type to explore the intricacies of interconnections between multiple layers of biological molecules and to identify system-level biomarkers^4–8^. Technology innovations and cost reduction have empowered increasingly large-scale multiomics studies for data collection on the same group of individuals, providing a revolutionary opportunity to fully understand and yield high-level insights into human diseases in a holistic fashion^9–14^.

Multiomics data integration can be classified into two categories depending on their objectives^15^. When the objective is on samples, the common multiomics integration strategy is data-driven clustering or classification of biological samples by combining complementary information contained in the multiomics data, followed by extracting system-level biologically differentiated networks for the endpoints such as wellness or disease subtyping^16–19^, or longitudinal trajectories^20–22^. When the objective is on the measured features, the data integration strategy is to identify significant multilayered molecular networks, so as to reveal the perturbed signatures and potential actionable targets for disease prevention and treatment^23–31^. Given the complexity of tying together multiomics data with unprecedented dimensionality and diversity, assigning accurate sample groups, and extracting true biological networks are challengeing^15, 32, 33^. Moreover, large-scale consortia based multiomics data are often generated across platforms, labs, and batches, creating unwanted variations and multiplying the complexities. Therefore, efficient data integration is essential for reliable multiomics studies^32^.

The data integration tasks in large-scale multiomics studies usually fall into two categories of application scenarios^34^. Horizontal (within-omics) integration, *i.e*., integration of diverse datasets from a single omics type, aims to combining multiple datasets across multiple batches, technologies, and labs from the same omics type for downstream analysis. Unwanted variations can result in systematic deviations (knowns as batch effects) confounded with critical study factors^35, 36^. Currently, various horizontal integration methods for bulk and single-cell omics data are available^37–39^. However, the selection of horizontal integration methods based on arbitrary visualizations of integrated datasets is challenging due to the lack of ground truth and objective quality control (QC) metrics for method selection. Vertical (cross-omics) integration, *i.e*., integration of diverse datasets from multiomics types, aims to combining multiple omics datasets with different modalities from the same set of samples, followed by designing appropriate downstream analysis to identify accurate sample groups, or multilayered and interconnected networks of biomolecular features^6, 34, 40, 41^.

Devising proper vertical integration strategies for sample clustering or feature identification is challenging in multiomics profiling. First, different technologies result in varying numbers of features and statistical properties, which can have a strong influence on the integration step to appropriately select and weigh different modalities. Secondly, each omics dataset has its intrinsic technological limits and noise structure. Combining multiomics datasets also multiply all the technical noises across different technologies, making it more challenging to integrate multiple datasets. Thirdly, many multiomics data integration algorithms and software are developed based on different statistical principles and assumptions^42–44^. Each multiomics integration method can report a solution, but assessing its reliability is difficult due to the lack of multiomics “ground truth” and QC methods for these complex processes.

Multiomics reference materials and relevant QC metrics are required for quality assessment of each omics measurement and its horizontal integration before successful multiomics-level vertical data integration^45–48^. Unrelated reference materials have been widely used as “ground truth” for performance evaluation of technologies for the same omics type, such as the genomic DNA^49,50^, tumor-normal paired DNA^51–53^, RNA, protein, or metabolite reference materials^54–57^. However, multiomics profiling requires measuring multiple types of omics data from the same set of interconnected reference samples, thus allowing for assessment of the ability to distinguish different reference samples with integrated datasets. Moreover, DNA, RNA, protein, and metabolite reference materials should be prepared simultaneously, which can provide “built-in truth” (the central dogma) for validating the hierarchical relationship among identified features. Therefore, publicly accessible and well-characterized multiomics reference materials at the genome scale are urgently needed^47^. Importantly, QC metrics relevant to research purposes are also critically important for assessing the quality of multiomics profiling. Precision and recall are widely used QC metrics for qualitative genomic variant calling^58–60^, whereas correlation coefficient is widely used for quantitative omics profling^55, 56, 61–64^. However, multiomics profiling is an integrated process, therefore the QC process should be performed based on the entire sample-to-result pipelines. Integrating multiomics information for more robust sample classifiers and multilayered interconnected molecular signatures are the major goals for multiomics profiling. Therefore, QC metrics should be related to these two research objectives, and should be suitable for evaluating the performance of each omics type ranging from data generation to multiomics data integration.

We launched the Quartet Project (chinese-quartet.org) to provide multiomics “ground truth” as well as best practices for the QC and data integration of multiomics profiling. The Quartet multiomics reference material suites, *i.e*., DNA, RNA, proteins, and metabolites developed from the B-lymphoblastoid cell lines derived from a quartet family of parents and monozygotic twin daughters, were designed for objectively evaluating the wet-lab proficiency in data generation and reliability of computational methods for horizontal data integration of the same omics type and for vertical data integration of multiomics types. A broad collection of the Quartet multiomics data generated from key technologies provides resources for evaluating the performance of new labs, platforms, protocols, and analytical tools. Based on the pedigree information of the Quartet samples, the horizontal and vertical data integration performance can be objectively evaluated, which provides unique insights into the commonly used multiomics integration strategies. We also developed a user-friendly data portal for the community to conveniently utilize and improve the Quartet resources (chinese-quartet.org). Most importantly, our study identifies “absolute” feature quantitation as the root cause of irreproducibility in multiomics measurement and data integration, and urges a paradigm shift from “absolute” to “ratio"-based quantitative multiomics profiling.

## Results

### Overview of the Quartet Project

The Quartet Project provides the community with multiomics reference materials and reference datasets for objectively assessing quality in data generation and integrated analysis in increasingly large-scale multiomics studies (**Fig. 1a**). Large quantities of multiomics reference materials suites (DNA, RNA, protein, and metabolite) were established simultaneously from the same immortalized B-lymphoblastoid cell lines (LCLs) of a Chinese Quartet family from the Fudan Taizhou Cohort^65^ (**Extended Data Fig. 1**), including father (F7), mother (M8), and monozygotic twin daughters (D5 and D6). As summarized in **Table 1**, each reference material was stocked in more than 1000 vials. They are suitable for a wide range of multiomics technologies, including DNA sequencing, DNA methylation, RNAseq, miRNAseq, LC-MS/MS based proteomics and metabolomics. Importantly, the DNA and RNA reference material suites have been approved by China’s State Administration for Market Regulation as the First Class of National Reference Materials and are extensively being utilized for proficiency testing and method validation. The Quartet multiomics design provides two types of “built-in truth” for quality assessment of multiomics profiling. One is the ability to correctly classify Quartet multiple samples, which is related to the multiomics research purpose of sample clustering. The metrics that measure the ability to correctly classify different Quartet samples are suitable for quality assessment of data generation and data analysis for each omics type. The other type of metric measures the ability to correctly identify the hierarchical relationships across multiomics features according to the rule of the central dogma, which can be used for assessing the reliability of correlation-based multiomics network integration.

**Table 1.** Summary of Quartet multiomics reference materials.

| Table 1 \| Summary of Quartet multiomics reference materials |  |  |  |  |  |  |  |  |
| --- | --- | --- | --- | --- | --- | --- | --- | --- |
| Type | Reference Material ID | Certified Reference Material ID | Amount | Cell Collection | Material Extraction | Description | User Scenarios | Quantity (Vials) |
| DNA | FDU_Quartet_DNA_D5_20160806 | GBW09900 | 10 µg | 20160806 | 20160819 | 220 ng/µL, 50 µL/tube in TE solution. DNA integrity number (DIN) > 8.5, with the peak size > 60,000 bp. | Short-read and long-read sequencing, microarray, PCR. | 1000 each |
|  | FDU_Quartet_DNA_D6_20160806 | GBW09901 |  |  |  |  |  |  |
|  | FDU_Quartet_DNA_F7_20160806 | GBW09902 |  |  |  |  |  |  |
|  | FDU_Quartet_DNA_M8_20160806 | GBW09903 |  |  |  |  |  |  |
|  | FDU_Quartet_DNA_D5_20171028 | / | 10 µg | 20171028 | 20171207 | 220 ng/µL, 50 µL/tube in TE solution. DNA integrity number (DIN) > 8.5, with the peak size > 60,000 bp. | Short-read and long-read sequencing, microarray, PCR. | 1000 each |
|  | FDU_Quartet_DNA_D6_20171028 | / |  |  |  |  |  |  |
|  | FDU_Quartet_DNA_F7_20171028 | / |  |  |  |  |  |  |
|  | FDU_Quartet_DNA_M8_20171028 | / |  |  |  |  |  |  |
| RNA | FDU_Quartet_RNA_D5_20171028 | GBW09904 | 5 µg | 20171028 | 20180725 | 520 ng/µL, 10 µL/tube in water solution. RNA integrity number (RIN) > 8.5. miRNA and other small RNA are retained. | RNA profiling, small RNA profiling. | 1000 each |
|  | FDU_Quartet_RNA_D6_20171028 | GBW09905 |  |  |  |  |  |  |
|  | FDU_Quartet_RNA_F7_20171028 | GBW09906 |  |  |  |  |  |  |
|  | FDU_Quartet_RNA_M8_20171028 | GBW09907 |  |  |  |  |  |  |
| Protein | FDU_Quartet_Pipetide_D5_20171028 | / | 10 µg | 20171028 | 20171106 | Dried, tryptic peptide mixtures. | Label-free LC-MS/MS-based proteomics. | 1000 each |
|  | FDU_Quartet_Pipetide_D6_20171028 | / |  |  |  |  |  |  |
|  | FDU_Quartet_Pipetide_F7_20171028 | / |  |  |  |  |  |  |
|  | FDU_Quartet_Pipetide_M8_20171028 | / |  |  |  |  |  |  |
|  | FDU_Quartet_Pipetide_D5_20171028 | / | 10 µg | 20171028 | 20200616 | Dried, tryptic peptide mixtures. Four labeled peptides are spiked in at different weight ratios as external controls. | Label-free LC-MS/MS-based proteomics. | 1000 each |
|  | FDU_Quartet_Pipetide_D6_20171028 | / |  |  |  |  |  |  |
|  | FDU_Quartet_Pipetide_F7_20171028 | / |  |  |  |  |  |  |
|  | FDU_Quartet_Pipetide_M8_20171028 | / |  |  |  |  |  |  |
| Metabolite | FDU_Quartet_Metabolite_D5_20171028 | / | 10<sup>6</sup> cells | 20171028 | 20200108 | Dried cell extracts from 1,000,000 cells using menthol: water solution. Ten external controls are spiked in at known amount. | LC-MS/MS-based metabolomics. | 1000 each |
|  | FDU_Quartet_Metabolite_D6_20171028 | / |  |  |  |  |  |  |
|  | FDU_Quartet_Metabolite_F7_20171028 | / |  |  |  |  |  |  |
|  | FDU_Quartet_Metabolite_M8_20171028 | / |  |  |  |  |  |  |

The Quartet multiomics reference materials were from monozygotic twin family, including father (F7), mother (M8), and monozygotic twin daughters (D5 and D6).

**Fig. 1.**
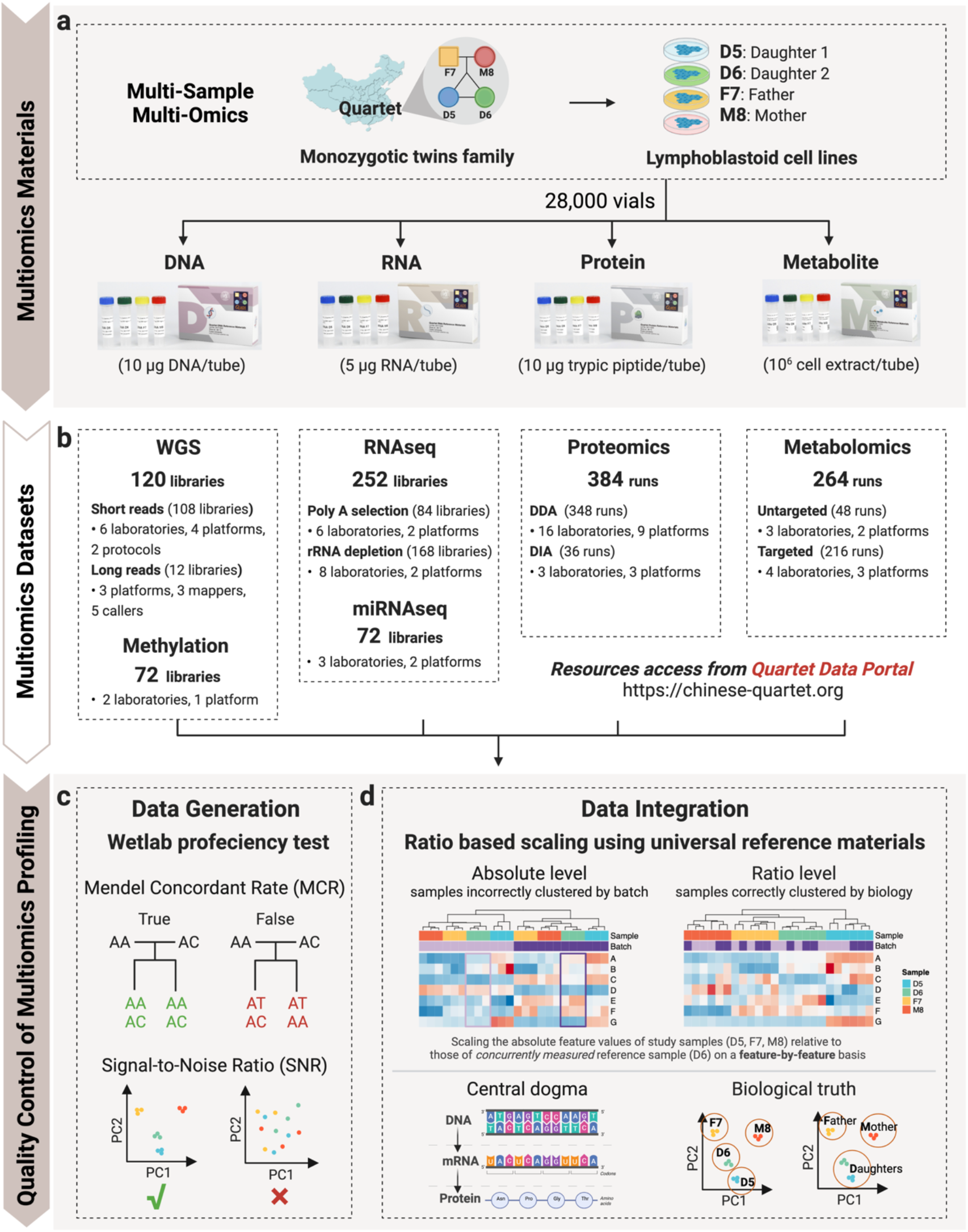
Overview of the Quartet Project. **a**, Design and production of Quartet family-based multiomics reference material suites. **b**, Data generation across multiple platforms, labs, batches, and omics types. **c**, Wet-lab proficiency test for the generation of each type of omic data using Quartet multi-sample-based reference materials. **d**, Ratio-based scaling using universal reference materials empower within-(horizontal) and cross-omics (vertical) data integration. Two types of QC metrics for multiomics data integration are developed: the cross-omics feature relationships that follow the central dogma, and the ability to classify samples into either four phenotypically different groups (D5-D6-F7-M8) or genetically driven three clusters (Daughters-Father-Mother).

The Quartet multiomics reference material suites were profiled across the commonly used multiomics platforms for comprehensive performance evaluation (**Fig. 1b**), including seven DNA sequencing platforms, one DNA methylation platform, two RNAseq platforms, two miRNAseq platforms, nine LC-MS/MS based proteomics platforms, and five LC-MS/MS based metabolomics platforms (**Extended Data Table 1**). Three technical replicates for each reference material were measured in each lab for performance evaluation, except for the long-reads DNA sequencing platforms where only one replicate was sequenced for each platform. **Extended Data Table 1** summarized the Quartet multiomics datasets for the real-world assessment of commonly used multiomics technologies. All the data can be accessed from the Quartet Data Portal (chinese-quartet.org), which provides a landscape of data quality for each type of omics profiling.

QC of multiomics profiling can be achieved using the Quartet multiomics reference materials. For data generation of each omics profiling, the Quartet built-in QC metrics, i.e., Mendelian concordance rate for genomic variant call and Signal-to-Noise Ratio (SNR) for quantitative omics profiling, enable proficiency testing on a whole-genome scale, which is essential for multiomics discovery studies (**Fig. 1c**). We proposed the ratio-based scaling using reference materials to empower horizontal and vertical omics data integration. The ratio-based data were derived by scaling the absolute feature values of study samples (D5, F7, and M8) relative to those of a concurrently measured reference sample (D6) on a feature-by-feature basis (**Fig. 1d**). In addition, the Quartet Project design provides two types of QC metrics to evaluate the reliability of vertical data integration for sample clustering. One QC metric is the cross-omics feature relationships that follow the rule of the central dogma. Another QC metric is to classify the Quartet samples correctly for both four different individuals (daughter1-daughter2-father-mother) and genetically driven three clusters (daughters-father-mother) (**Fig. 1d**). A reliable vertical integration method needs to be able to discover the intricate biological differences under these scenarios.

In this article, we described the overview of the Quartet Project, including the performance of multiomics technologies and data integration strategies, and the best practice guidelines for process control of large-scale multiomics studies using the Quartet reference materials (**Extended Data Fig. 2**). Four accompanying papers detailed the establishment of the DNA^66^, RNA^67^, protein^68^, and metabolite^69^ reference materials, reference datasets, and QC methods for each type of omics profiling (genomics, transcriptomics, proteomics, and metabolomics, respectively) and their applications. Another paper^70^ was dedicated to addressing the widespread problem of batch effects present in each and every type of omics data. We also developed the Quartet Data Portal (chinese-quartet.org)^71^ for the community to conveniently access and share the Quartet multiomics resources according to the regulation of the Human Genetic Resources Administration of China (HGRAC).

**Fig. 2.**
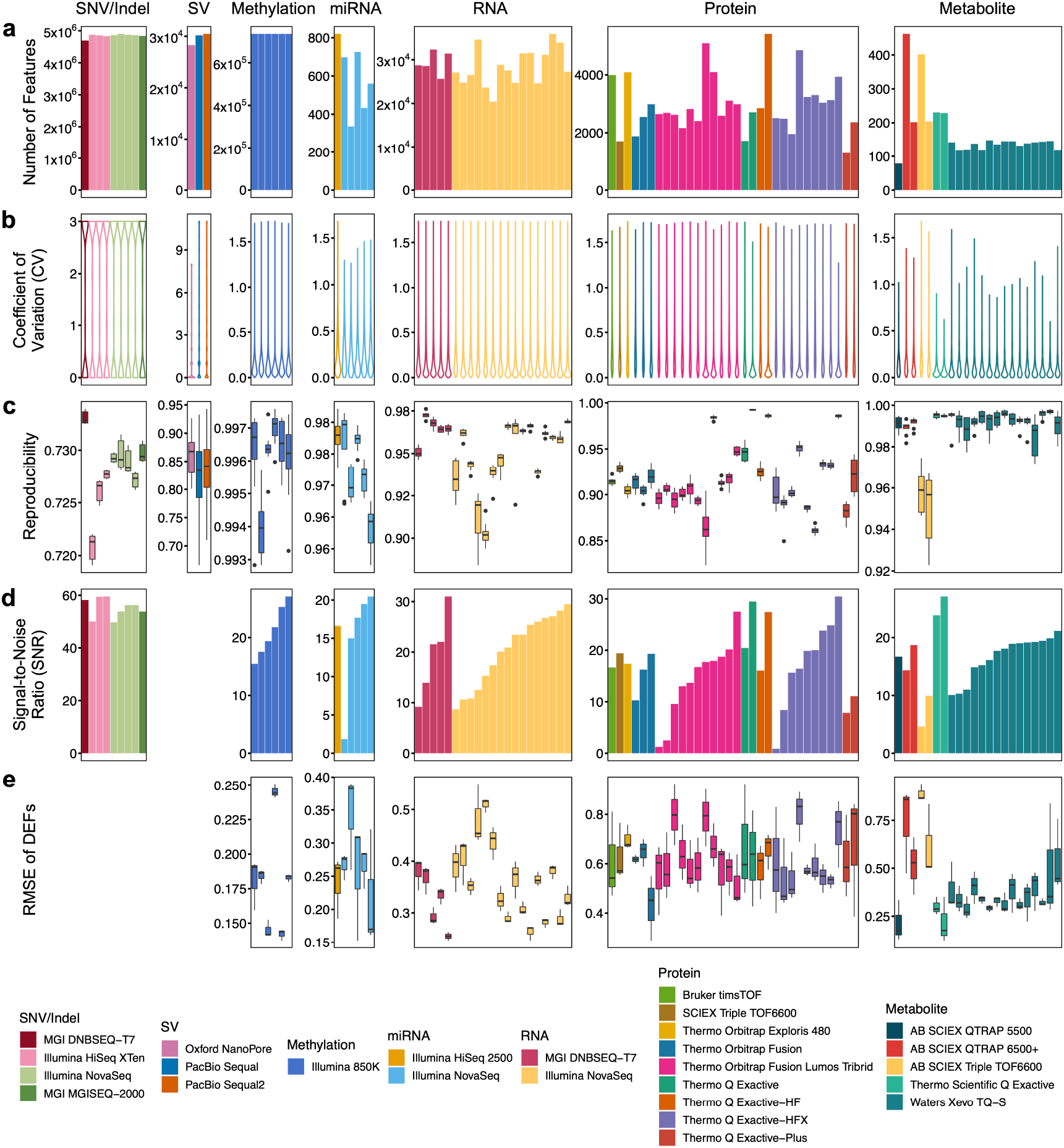
Wet-lab proficiency diverges in multiomics data generation. **a**, The number of features detected from each dataset generated in different labs using different platforms. **b**, Distribution of the number of experiments supporting genomic variant calling, or coefficient of variation (CV) in quantitative omic profiling from technical replicates (analytical repeats in SV calling, and library repeats for the others) within a batch. **c**, Technical reproducibility from three replicates within a batch, calculated as the Jaccard index for genomic variant calling and Pearson correlation coefficient (*r*) for quantitative omic profiling. **d**, Signal-to-Noise Ratio (SNR) based on Quartet multi-sample design (4 samples × 3 replicates per batch). **e**, Root Mean Square Error (RMSE) of high confidence differentially expressed features (DEFs).

### Wet-lab proficiency in data generation varies substantially within every omics type

The performance of each omics platform in different labs was assessed using the Quartet multiomics reference materials. Except for the long-reads sequencing platforms, the reference materials were profiled within a batch in a lab (4 samples × 3 replicates). For long-reads sequencing, one replicate for each reference material was sequenced, and the resulting data were analyzed using 11 pipelines, therefore the performance evaluation was conducted only at the analytical procedure level. Details on data generation and analysis were given in the Methods section.

QC metrics for objective performance evaluation are critically important. The number of measured features, coefficient of variation (CV), and technical reproducibility are widely used QC metrics across different omics platforms, and were used in our study for cross-omics performance comparisons. As shown in **Fig. 2**, the number of features measured by each omics type varied by several orders of magnitude, from 60 metabolites to 4.8 million DNA small variants (SNVs and Indels) (**Fig. 2a**) per sample. Within each omics type, the number of features detected varied among batches and labs. There were no obvious differences in the numbers of detected small variants between Illumina and BGI platforms, with each platform detecting approximately 4.8 million small variants. However, the numbers of detected proteins among different LC-MS/MS based proteomics platforms were more profound. The reproducibility of detected features in each omics profiling was evaluated using the number of replicates supporting a variant call for genomics and the coefficient of variation (CV) in quantitative omics profiling among technical replicates within a batch (**Fig. 2b**). Most SNVs were supported by all the three library replicates within the batch (Jaccard index of ~0.94 for SNVs), whereas the number of analytical repeats supporting a structural variant (SV) call greatly varied (Jaccard index of ~0.70 for SVs), indicating large differences in SV calling among analytical pipelines.

For quantitative omics profiling, the CVs of most quantified features were below 30%. In addition, the reproducibility of technical replicates was also evaluated at individual sample level (**Fig. 2c**). Reproducibility was calculated as the Jaccard index from three library repeats within a batch. For the short-reads sequencing platforms, all Jaccard index values were above 93%. Moreover, the reproducibility of SV from 11 call sets using different analytical pipelines was between 80% and 90%. Nanopore was found to be more reproducible than PacBio among the long-read sequencing platforms. The reproducibility of quantitative omics profiling was calculated as the Pearson correlation coefficient (Pearson *r*) of technical replicates within a batch. The *r* values from all labs and metabolomic platforms were above 95%, indicating high reproducible in metabolomic profiling for the same sample. However, the *r* values for repeated measurements of the same sample were between 88.42% and 97.62% for transcriptomics, or from 82.37% to 99.34% for proteomics (**Fig. 2c**).

Based on the Quartet multi-sample design, we defined two QC metrics to measure the ability to identify intrinsic biological differences among various groups of samples, a key objective of omics profiling. The Quartet based Signal-to-Noise Ratio (SNR) is the ratio of inter-sample differences (*i.e*., “signal”) to intra-sample differences among technical replicates *(i.e*., “noise”). It is calculated as the ratio of the average distance between the Quartet samples to the average distance between technical replicates of the same sample (see Methods for details). For a measurement method with high resolution in differentiating biologically different groups of samples, the inter-sample differences of Quartet samples should be much larger than the variation among technical replicates of the same sample. Principal component analysis (PCA) showed clear separation among the Quartet samples (D5, D6, F7, and M8) for high-quality profiling experiments (**Extended Data Fig. 3a**) but not for low-quality profiling experiments (**Extended Data Fig. 3b**). Strikingly, high variabilities in intra-batch data quality were observed in each omics platform (**Fig. 2d**), especially for the quantitative omics platforms, such as transcriptomics (SNR range 8.7 – 31.0, SD=7.1), miRNA profiling (SNR range 1.9 – 20.5, SD=6.8), proteomics (SNR range 0.9 – 30.5, SD=7.5), and metabolomics (SNR range 4.6 – 27.1, SD=5.1). Moreover, variabilities within the same technology platform were higher compared to those between different platforms of the same omics type. For example, both high and low SNRs were observed in RNAseq for the Illumina and BGI platforms, but the average SNRs of the two sequencing platforms were very close (20.39 vs. 19.54, *p*=0.84). Similarly, high variabilities in SNR were observed within each MS platform for proteomics or metabolomics profiling. These results implied that proficiency of an individual wet-lab, instead of a specific platform itself, was a more important factor affecting the reliability of data generation for each omics type.

**Fig. 3.**
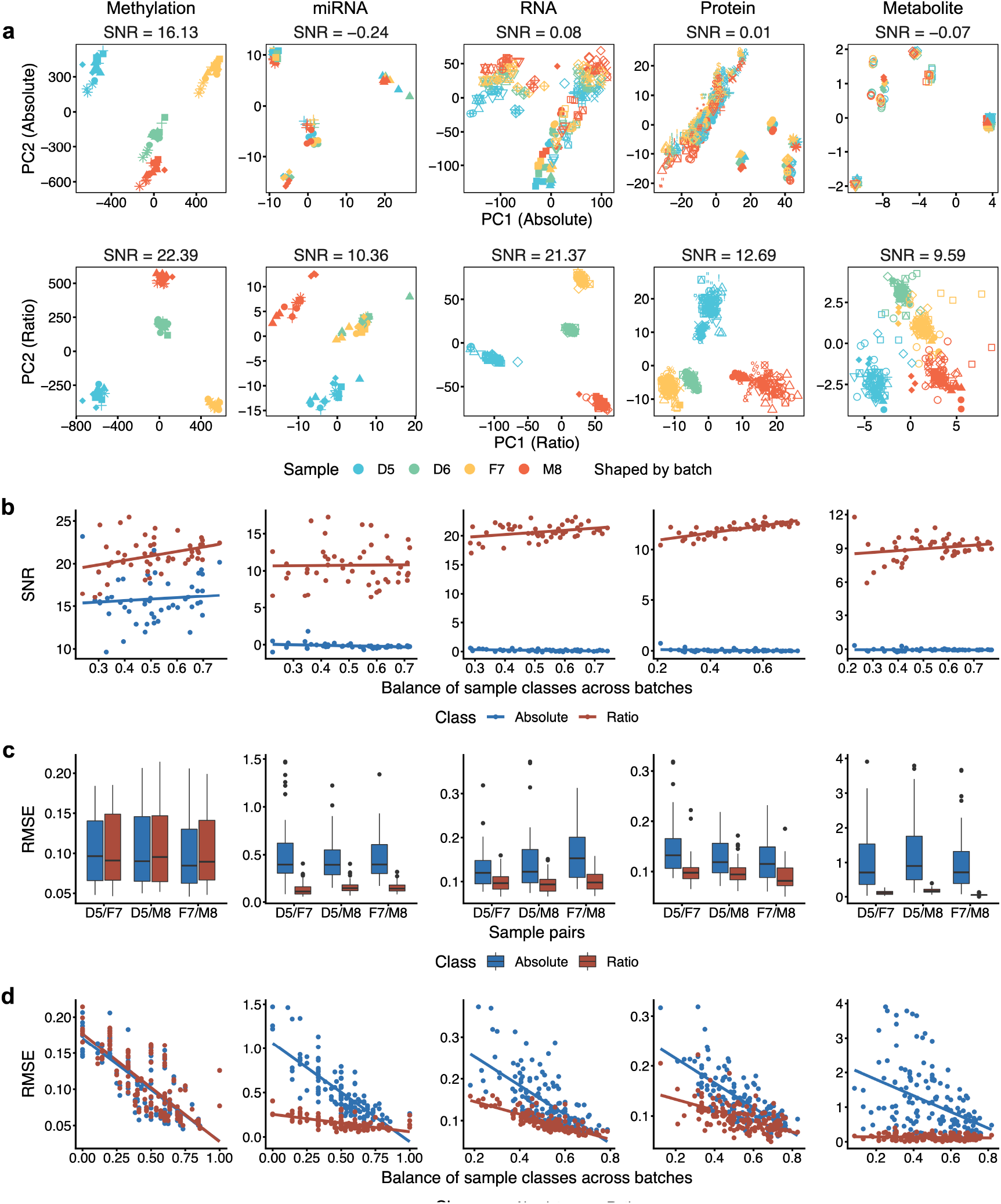
Ratio-based scaling enables horizontal integration of datasets across batches, labs, and platforms. **a**, PCA plots of horizontal integration of all batches of methylation, miRNAseq, RNAseq, proteomics, and metabolomics datasets at absolute level (raw data, top row) and ratio level (ratio scaling to D6 sample, bottom row). **b**, Scatter plots between SNR and degree of sample class-batch balance. Blue: absolute level; Red: ratio level. **c**, Boxplots of RMSE of the DEFs of horizontal integration data at absolute level (Blue) and ratio level (Red) based on the reference datasets. **d**, Scatter plots between RMSE when integrating at absolute (Blue) and ratio (Red) levels and the degrees of sample class-batch balance.

In addition, we constructed high-confidence reference datasets of differentially expressed features (DEFs) in terms of the level of differential expression between a pair of samples (D5/F7, D5/M8, and F7/M8) for each quantitative omics profiling using a consensus-based integration strategy (**Extended Data Fig. 4**). Therefore, the Root Mean Square Error (RMSE) was used for quantitatively evaluating the consistency between a test dataset with the high-confidence reference dataset (**Fig. 2e**). One major goal of each omics profiling is to identify molecular features that are intrinsically different between distinct sample groups such as disease versus normal, or responders versus non-responders to a drug treatment. Thus, the ability to accurately differentiate biologically different groups of samples is a critical metric for measuring the performance of a technology, procedure, or lab.

**Fig. 4.**
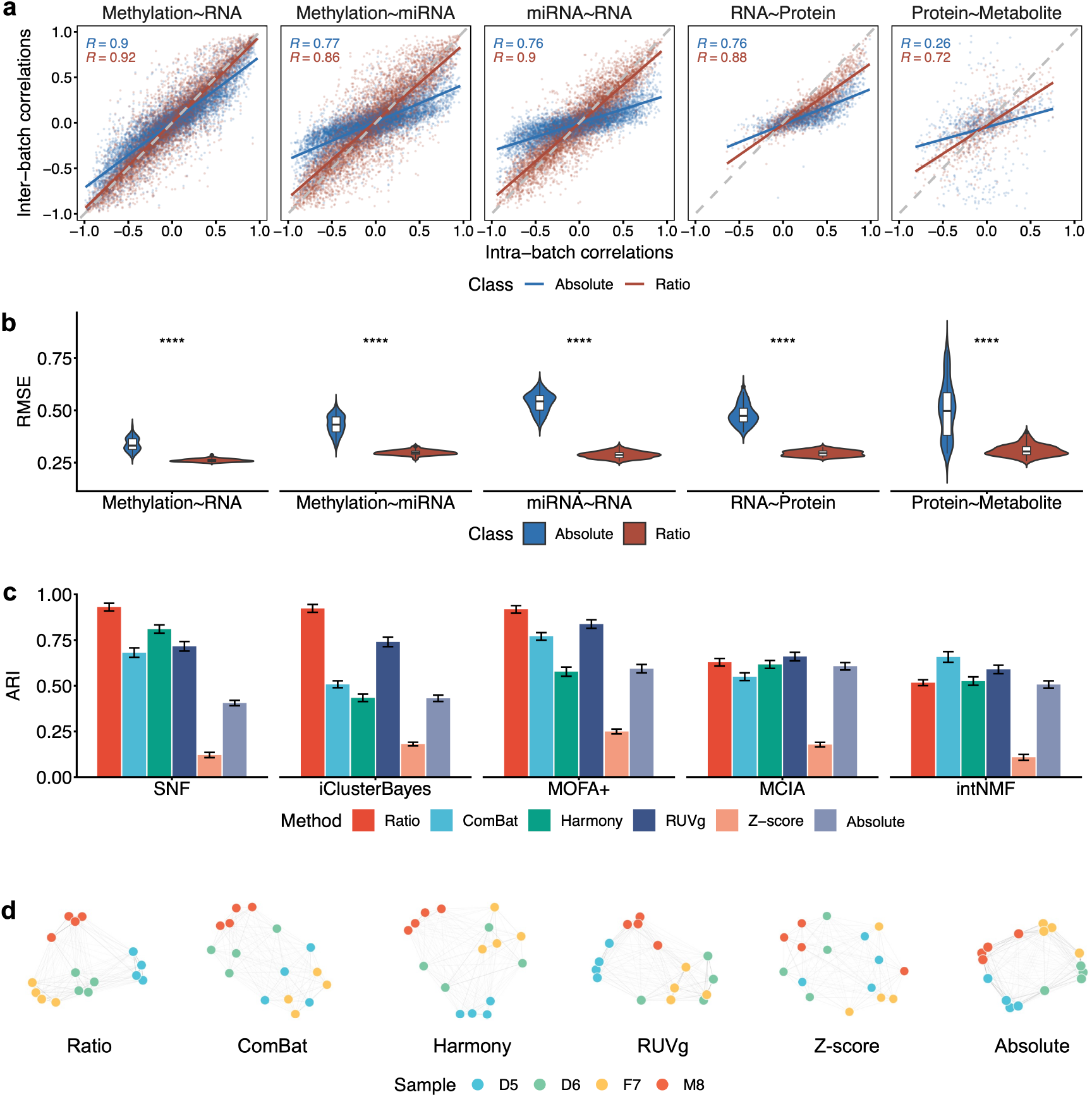
Ratio-based scaling facilitates vertical integration of datasets from different omics types. **a**, Scatter plots between cross-omics feature relationships of intra- and inter-batch (horizontally integrated) data at absolute level (Blue) and ratio level (Red). The solid lines represent fitted curves from linear regression along with the Pearson correlation coefficients. **b**, Violin plots of RMSE of cross-omics feature relationships of horizontal integration data at absolute level (Blue) and ratio level (Red) based on the reference datasets. **c**, Bar plots of the Adjusted Rand Index (ARI) of vertically integrated multiomics datasets of multiple batches using different algorithms including SNF, iClusterBayes, MOFA+, MCIA, and intNMF. Data of each omics type were preprocessed by Ratio, ComBat, Harmony, RUVg, Z-score, or Absolute (no further processing on the normalized datasets) for horizontal integration. **d**, Sample similarity networks for SNF integration with different data preprocessing methods in **c**.

We explored the relationships between SNR and the number of detected features, the reproducibility of features, the reproducibility of technical replicates, and the RMSE of DEFs for identifying quality issues in quantitative omics profiling (**Extended Data Fig. 5**). These data suggested that high correlation coefficients between repeated measures of the same sample did not guarantee high resolution (SNR) in identifying inherent differences (*i.e*., biological signals) among various biological sample groups. Therefore, multi-sample-based QC metrics are needed in identifying labs with low proficiency in detecting intrinsic biological differences among sample groups.

**Fig. 5.**
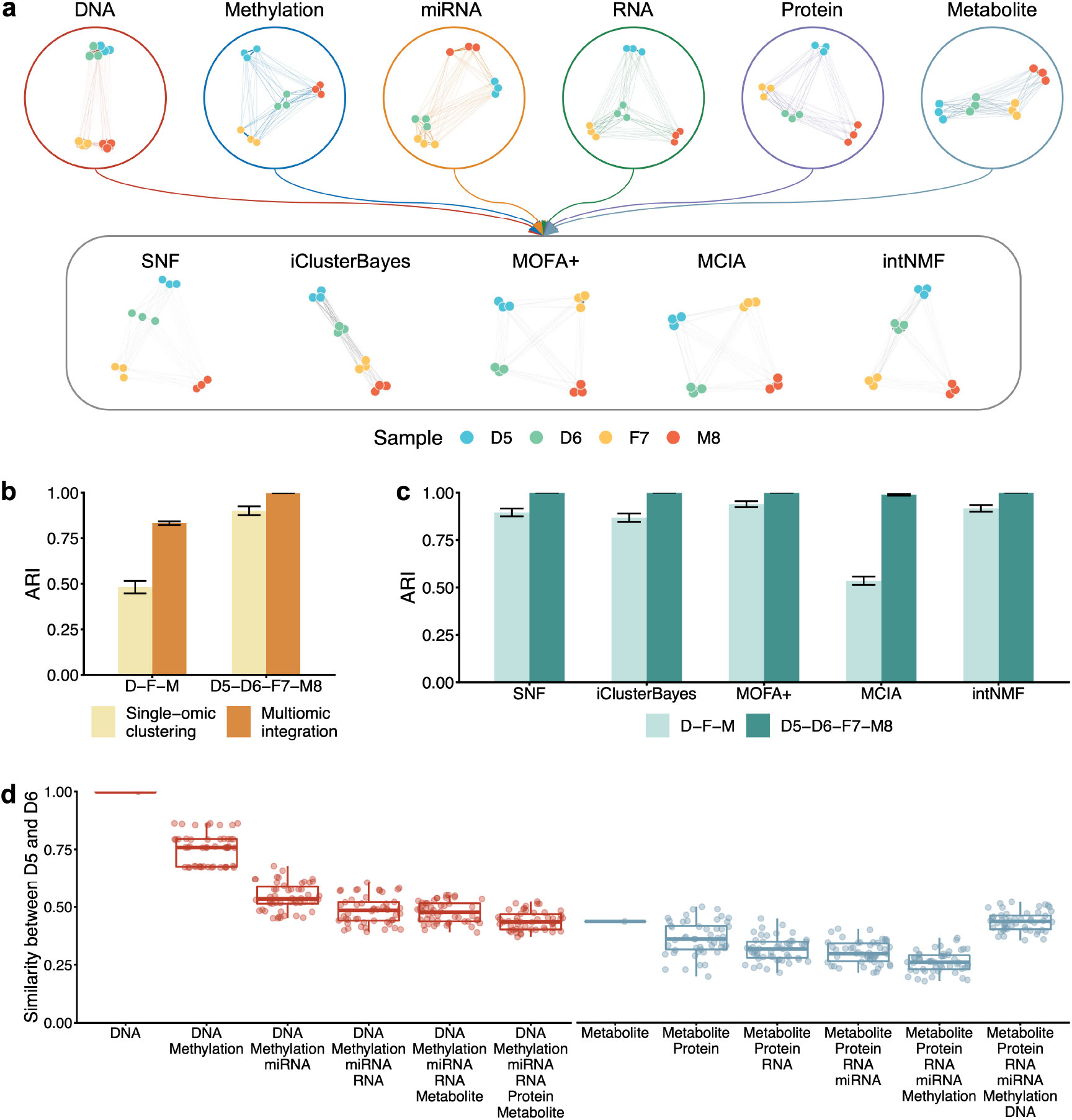
Quartet multiomics design provides genetics-driven ground truth for vertical integration. **a**, Networks of six types of omics based on the similarity between 12 samples within one batch (top row), and sample similarity networks obtained by SNF, iClusterBayes, MOFA+, MCIA, and intNMF (bottom row) that integrated the six types of multiomics data. **b**, Bar plots of the ARIs when clustering samples into three (D-F-M) or four groups (D5-D6-F7-M8) by single-omic (Light yellow) versus multiomics integration (Dark yellow) using PAM clustering algorithms. **c**, Bar plots of the ARIs of multiomics data integration using SNF, iClusterBayes, MOFA+, MCIA, and intNMF. Light green represents data when the true labels of the samples were set to three clusters (D-F-M), and dark green represents four clusters (D5-D6-F7-M8). **d**, Box plots of the ARIs for integration of different types of omic data. The multiomics data were integrated started from DNA (Red), and metabolite (Grey) by using SNF.

### Ratio-based scaling enables horizontal integration of data across platforms, labs, and batches for the same omics type

In large-scale omics studies, reliability of horizontal integration of omics datasets across different platforms, labs, or batches for the same omics type is even more challenging compared to within-batch proficiency mentioned above. If the Quartet reference materials were profiled per-batch along with study samples in each lab, the reliability of horizontal integration could be assessed by the Quartet multi-sample based SNR. Horizontally integrated datasets should have the ability to differentiate the Quartet samples. In order to evaluate the reliability of horizontal integration of each omics profiling in differentiating the four sample groups in the Quartet design, we integrated datasets for all batches of the same omics type separately, including methylation array data (M value), miRNAseq (log2CPM), RNAseq (log2FPKM), proteomics (log2FOT), and metabolomics (log2Intensity) (**Fig. 3**).

Horizontal integration of the aforementioned five types of quantitative omics data all showed obvious batch-dominant clustering at absolute expression levels (**Fig. 3a** top). However, after converting the absolute omics data to a ratio scale relative to the same reference material (D6) within a batch on a feature-by-feature basis, PCA plots showed clear separation of the four types of reference samples (D5, D6, F7, and M8) and the drastic batch effects seen at the absolute scale largely disappeared (**Fig. 3a** bottom). We further quantitatively measured the quality of horizontal data integration using the Quartet multi-sample based SNR as the metric. A method of good quality for horizontal data integration at each omics level would clearly separate the four Quartet sample groups, *i.e*., the inter-sample differences of the Quartet samples should be much larger than the variation among technical replicates. As shown in **Fig. 3a**, the SNR after horizontal integration of datasets for each omics type at the absolute level was all close to zero except for methylation data (**Fig. 3a** top), whereas the SNR of the integrated datasets dramatically increased at the ratio level (**Fig. 3a** bottom). Importantly, these conclusions remain the same if one chooses D5, F7, or M8 instead of D6 as the reference sample (**Extended Data Fig. 6**), indicating the universal applicability of the ratio-based scaling approach.

In addition, we characterized the impact of the level of batch effect on horizontal-integration SNR by randomly selecting samples from different batches and using the average of the Jaccard index of the batches from the four sample groups as a measure of group-batch balance. As shown in **Fig. 3b**, regardless of the level of balance of sample classes across batches, horizontal integration at the ratio level resulted in much better discrimination between sample classes, *i.e*., much higher SNR. However, the corresponding SNR at the absolute level was all close to zero except for methylation data, whether there was a group-batch balance or not. These results clearly demonstrated that quantitative omics profiling at the ratio level was much more comparable and suitable for horizontal integration than at the absolute level.

Ratio-based profiling allows for more accurate determination of the subtle differences between two Quartet samples on a feature-by-feature basis. For all three comparisons (D5/F7, D5/M8, and F7/M8), compared to the log2 fold differences of the absolute-based integration data, those of the ratio-based integration data showed a higher level of agreement (and lower RMSE) with the corresponding reference dataset for each omics type (**Figs. 3c** and **3d**). Furthermore, the level of balance of sample groups between batches was helpful for the accurate detection of DEFs. This was reflected in the negative correlation between RMSE and the level of group-batch balance (**Fig. 3d**). It was also clear that the lack of group-batch balance affected absolute-based data integration much more severely than ratio-based data integration, where the former showed a much larger slope than the latter (**Fig. 3d**).

The pervasiveness of batch effects in quantitative analysis techniques at the absolute expression level presents a real challenge for horizontal integration. Our results demonstrated that the conversion of quantitative omics data to a ratio scale relative to a common reference sample (*e.g*., the Quartet D6 sample) can effectively mitigate the detrimental impact of batch effects on sample classification, differential feature identification, etc.

### Ratio-based scaling facilitates vertical integration of data from different omics types

In large multiomics studies, the multiomics datasets are usually generated in multiple batches, platforms, and labs. Vertical integration of multiomics datasets from various omics types is typically performed after horizontal integration at the same omics type, thus the final integration performance is influenced by both horizontal and vertical dimensions. Therefore, we evaluated the reliability of vertical integration of horizontally integrated ratio-based data under different scenarios.

One advantage of multiomics studies is to systematically discover the cross-omics relationships from multiple interconnected biological layers. The Quartet multiomics design can provide the “built-in truth” from the hierarchical relationships across omics features. Since the Quartet multiomics reference materials were derived from the same batch of cultured cells, the generated multiomics data can be used to evaluate their compliance with the central dogma principle, *i.e*., how the genetic information is transcribed from DNA to RNA, and then translated to protein. The central dogma-regulated relationships between cross-omics features can be measured using the correlation coefficient, which is one of the simplest ways to estimate the pairwise relevance between two types of omics features, forming the basis of multiomics integration for network analysis.

Cross-omics feature relationships calculated based on multiple batches of data integrated at the ratio level (Inter-batch) showed stronger correlations with the cross-omics single batches (Intra-batch) than that at the absolute level (**Fig. 4a**). These cross-feature correlations of methylation-miRNA, methylation-RNA, miRNA-RNA, RNA-protein, and protein-metabolite were derived from features of both omics types associated with the same genes, which may more closely follow the principle of the central dogma. In particular, for the relationships between proteins and metabolites, direct integration of multi-batch data at the absolute level could not easily identify the true correlations between cross-omics feature pairs.

To more accurately measure the effect of integration based on ratio profiling, we constructed the Quartet reference datasets of the Pearson correlation coefficients between the expression levels of two different types of omics features in order to evaluate the performance of vertical integration at the feature relationship levels (**Extended Data Fig. 7**). The central dogma was reflected in the Quartet multiomics data as the abundance of RNAs was almost exclusively positively correlated with that of proteins in the reference dataset (224 RNA-protein pairs were positively correlated and no RNA-protein pair was negatively correlated). In agreement with **Fig. 4a**, the concordance of correlation coefficients of cross-omics features with the reference Pearson *r* was higher (as indicated by lower RMSEs) in the horizontally integrated data based on the ratio level compared to absolute level (**Fig. 4b**).

Another advantage of vertical integration of multiomics data is to be able to distinguish subtypes of clinical samples with subtle differences that cannot be identified based on a single type of omics data. Therefore, the ability to discover the true biological differences between sample groups is a key metric to measure the performance of multiomics integration tools and procedures. The multi-sample and multiomics design of the Quartet Project provides unique “ground truth” for assessing the reliability of vertical integration. Here we included six horizontal integration methods for evaluation, *i.e*., ratio-based scaling (Ratio), ComBat^72^, Harmony^73^, RUVg^74^, Z-score, and direct integration of the normalized values (Absolute). Five widely accepted vertical-integration tools were subsequently used, *i.e*., SNF^5^, iClusterBayes^75^, MOFA+^76^, MCIA^77^, and intNMF^78^, generating 30 combinations of horizontal and vertical integration for performance assessment.

The Adjusted Rand Index (ARI)^79^ is a widely used QC metric to compare clustering results against external criteria. To quantitatively evaluate the reliability of vertical data integration at the multiomics level, we used ground truth-based ARI (daughter1-daughter2-father-mother, *i.e*., D5-D6-F7-M8 as four independent sample groups or clusters) as the metric.

Ratio-based scaling data largely outperformed other horizontal-integration methods with a much higher ARI when the same vertical-integration algorithm was used (**Fig. 4c**). The final performance of integration was influenced by both horizontal-integration methods and vertical-integration algorithms. For example, regardless of which horizontal integration method was used, SNF performed better overall than intNMF in subsequent vertical integration. Furthermore, the multiomics-based sample similarity networks constructed based on the SNF clearly demonstrated the different power of correctly clustering the four Quartet sample groups by the six horizontal-integration methods (**Fig. 4d**). Integration using ratio-based profiling data showed tighter connections between objects with the same sample group (same color) and looser connections between objects from different groups. These results implicated that ratio-based scaling improved the vertical integration of sample clusters through reliable cross-sectional integration.

### Quartet multiomics design provides genetics-driven ground truth for vertical data integration

In addition to the relatively simple task of differentiating the four different individuals (daughter1-daughter2-father-mother, D5-D6-F7-M8), the Quartet monozygotic twin family design also provided a more challenging task of classifying these samples into the Quartet family-based and genetically distinct three groups (daughters-father-mother, D-F-M). Here we integrated the multiomics data of moderate quality (SNR in the range of top 20% to 80%) including DNA, methylation, miRNA, RNA, protein, and metabolite. For each vertical integration method, only one batch of data was selected for each omics type to prevent the influence of batch effect in horizontal integration. In addition, we conducted the Partitioning Around Medoids (PAM) clustering^80^ for each type of single omics data and calculated ARI as a control to assist in assessing the performance of the vertical integration.

The inter-sample similarity networks built using data from a single omics (top) and integrated multiomics data using SNF, iClusterBayes, MOFA+, MCIA, and intNMF (bottom) were visualized in **Fig. 5a**. At the DNA level, identical twin samples (D5 and D6) were tightly clustered together due to their near-identical DNA sequences. On the other hand, they showed no clear tendency of clustering together for all five types of quantitative omics data (methylation, miRNA, RNA, protein, and metabolite), and may even look relatively far apart (e.g., D6 and F7 appeared closer in miRNA, RNA, or protein data). This distinction in clustering tendency between DNA variants and quantitative omics data implied that the classification task (D-F-M) can be used to assess whether a vertical integration approach can reveal the intrinsic built-in genetic truth in the Quartet identical twin family.

Vertical integration reduced technical noise and improved the ability of sample clustering, indicated by the fact that the ARIs of both three clusters (D-F-M) and four clusters (D5-D6-F7-M8) of multiomics integration were higher than the direct clustering of single-omics data (**Fig. 5b**). Nevertheless, there were still differences in performance between the integration algorithms when distinguishing the three sample categories (D-F-M). SNF, iClusterBayes, MOFA+, and intNMF correctly classified these samples into three Quartet family-based groups (D-F-M), whereas MCIA did not perform well (**Fig. 5c**). This demonstrated that the integration algorithms could be prioritized by whether they found potential genetic truth (identical twins) behind the four individuals with distinct differences in molecular phenotypic data.

The similarity between identical twins (D5 and D6) during the vertical integration can be quantified to illustrate the impact of adding different layers of omics information on the clustering of Quartet samples (Methods for the details). As shown in **Fig. 5d**, the similarity between D5 and D6 decreased both when gradually adding downstream omics data starting from genomics, and when integrating upstream omics data starting from metabolomics (except for the eventual addition of DNA). This phenomenon again demonstrated that the genetic relationships between the Quartet identical twins were only reflected at the DNA level, and it also specified the need to incorporate genomic data when using the three clusters (D-F-M) as a QC metric for vertical integration.

### Best practice guidelines for QC and data integration of multiomics profiling using the Quartet reference materials

QC is comprised of procedures to ensure the reliability of multiomics profiling using defined QC metrics and thresholds to meet the requirements of different research purposes. Large-scale multiomics studies involve multi-center and long-term measurements where unified QC metrics and universal integration strategies are needed to ensure quality during data generation and integration. We recommend including the Quartet reference materials (*e.g*., four samples × three replicates) when profiling each batch of study samples, and propose the best practice guidelines for QC and data integration in the three aspects including intra-batch data generation, horizontal integration, and vertical integration (**Table 2**).

**Table 2.** Best practice guidelines for quality control and data integration of multiomics profiling using the Quartet reference materials.

| Procedures | Recommendations |
| --- | --- |
| **Data generation** | **QC metrics**<br>• Reference-free: Signal-to-Noise Ratio (SNR) for measuring the ability to differentiate biologically different Quartet samples; Mendelian concordance rate (MCR) for genomic data.<br>• Reference-based: precision, recall and F1-score for qualitative values; Root Mean Square Error (RMSE) for quantitative values. |
|  | **Thresholds**<br>• Minimum requirement for a reliable measurement is to differentiate the inherent differences between different sample groups.<br>• Relative quality ranking among accumulating datasets can be obtained through the Quartet Data Portal. |
|  | **Implementation**<br>• Perform a full performance validation for new labs, platforms, tests, or analytical pipelines.<br>• Participate in proficiency tests for independent assessment of test performance through inter-lab comparison. |
| **Horizontal (Within-omics) integration** | **QC metrics**<br>• Reference-free: Signal-to-Noise Ratio (SNR) for measuring the ability to differentiate biologically different Quartet samples; Mendelian concordance rate (MCR) for genomic data.<br>• Reference-based: precision, recall and F1-score for qualitative values; Root Mean Square Error (RMSE) for quantitative values. |
|  | **Thresholds**<br>• As long as the horizontally integrated datasets still have the ability to differentiate the different Quartet sample groups, the reliability of the follow-up exploratory studies is assured. |
|  | **Implementation**<br>• Profile the universal Quartet samples each batch along with study samples.<br>• Paradigm shift from “absolute” to “ratio”-based scaling by incorporating universal reference materials, which is inherently reproducible and batch-effect resistant. |
| **Vertical (Cross-omics) integration** | **QC metrics**<br>• Quartet family based: Adjusted Rand Index to classify samples into four clusters (D5-D6-F7-M8) and three clusters (Daughters-Father-Mother). Vertical integration should have the ability to discover Quartet family-based sample clustering (Daughters-Father-Mother).<br>• Central dogma based: built-in interconnection across omics features. Root Mean Square Error (RMSE) based on reference datasets. |
|  | **Thresholds**<br>• ARI ranges from 0 to 1, the larger the better. |
|  | **Implementation**<br>• Profile Quartet samples for each omic type along with study samples.<br>• Paradigm shift from “absolute” to “ratio”-based scaling by incorporating universal reference materials, which empowers vertical data integration. |

We provided both reference dataset-free and reference dataset-based QC metrics to assess wet-lab proficiency of data generation for the same omics type in terms of the capability of identifying the subtle differences between sample groups. Without relying on the reference datasets, the Quartet-based SNR (D5-D6-F7-M8) can be calculated for quality assessment for all types of omics data. The SNR calculated based on the four Quartet sample groups was more sensitive for assessing wet-lab proficiency than generic QC metrics based on multiple technical replicates of a single sample (**Fig. 1**). We also recommend the use of Mendelian concordance rate (MCR) based on the pedigree of the Quartet as a QC metric for assessing the quality of genomic data^66^. With the reference datasets, the wet-lab proficiency was assessed by the concordance between the evaluated batch of data and the reference datasets. Precision, recall, and F1-score were recommended for qualitative omics (small variants and structural variants), and RMSE at the ratio level (scaling to D6) of the feature expressions and the differential expressions between groups (D5/F7, F7/M8, and M8/D5) were recommended for quantitative omics (DNA methylation, transcriptomics, proteomics, and metabolomics). In addition, more comprehensive proficiency tests or inter-lab comparisons can be performed by obtaining the relative quality ranking among the cumulative datasets within the Quartet Data Portal^71^.

For horizontal integration of multi-batch data, a paradigm shift from “absolute” to “ratio"-based profiling by incorporating universal reference materials is essential and improves the reproducibility and batch-effect resistance. QC metrics used in intra-batch data generation can still be used in the quality assessment of horizontal integration. The reliability of further exploratory studies can be ensured as long as the horizontally integrated dataset can still distinguish different Quartet samples.

Vertical integration can be enhanced by ratio scaling the data based on reference materials. The Quartet multiomics and multi-sample reference materials provide two types of “built-in truth” for QC of vertical integration. The first type of “built-in truth” is based on the clustering of Quartet samples through the combined use of ARI_D-F-M_ and ARI_D6-D6-F7-M8_ to synthetically characterize the quality of vertical integration. In addition, the ability to correctly distinguish samples into four clusters (D5, D6, F7, and M8), as measured by ARI_D6-D6-F7-M8_, indicates that the integrated multiomics data must have the basic ability to differentiate the four different biological samples from technical replicates. On the other hand, the integration algorithm must be able to identify the multiomics features driven by the built-in genetic truth of the Quartet identical twin, thus separating samples into three clusters (daughters D, father F, and mother M) by identifying true cross-omics associations. The second type of “built-in truth” is the hierarchical relationship across omics features following the principle of the central dogma. RMSE of cross-omics feature relationships calculated based on the reference datasets can be used to evaluate the accuracy of the cross-omics feature correlations.

## Discussion

We developed the first suites of publicly available multiomics reference materials, including matched DNA, RNA, protein, and metabolite from immortalized B-lymphoblastoid cell lines of four individuals of a Chinese quartet family. We then extensively profiled these reference materials using diverse multiomics technology platforms in multiple labs across batches with repeated measurements. The reference datasets of measurands characterizing these reference materials at the genome scales were established based on a consensus approach using multiple bioinformatics pipelines and data integration approaches. The reference materials and the reference datasets can facilitate objective quality assessment of multiomics profiling (**Table 1**) by providing two types of “built-in truth” for QC of multiomics data generation and data integration. One is about the clustering of the Quartet samples based on their intrinsic biological differences, and the other is about the inherent relationships across omics features following the central dogma’s rule (DNA to RNA to protein). The resulting wealth of multiomics resources were made publicly available via the Quartet Data Portal (chinese-quartet.org).

Wet-lab proficiency was consistently found to be a more important factor affecting the quality of data generated for each omics type than the choice of a specific technology platform (**Fig. 2**). Our findings are consistent with what have been reported previously on gene-expression profiling with microarrays in MAQC-I^55^ and with RNAseq in MAQC-III (SEQC)^54^ when the same pair of MAQC reference RNA samples A (a mixture RNA of ten cancer cell lines) and B (a mixture RNA from brain tissues of 23 donors) were analyzed by a given platform in multiple labs. This observation seems intuitive; however, no adequate solution has been validated or adopted by the scientific community, hence has likely contributed to the lack of reproducibility of biomedical research^81^. Our observation highlights the urgency of highly sensitive proficiency testing to improve internal lab proficiency before profiling precious research and clinical samples. To this end, we established appropriate reference materials and proposed sensitive metrics for performance assessment.

The ability to correctly identify molecular phenotypic differences between various groups of samples or clinical subtypes of a disease is a fundamental requirement for any omics technology-based research. Thus, an appropriate performance metric should be taken into account and multiple groups of samples must be included to meet this vital requirement. For each omics type, the Quartet study design included four groups of samples (D5-D6-F7-M8), allowing us to define the universal Signal-to-Noise Ratio (SNR) metric for measuring the performance of any multiomics technologies. We found that the SNR metric was sensitive in identifying low-quality datasets that may otherwise be considered as of high quality. For example, reproducibility of repeated measurements (or technical replicates) of the same sample, usually expressed as coefficient of variation, Pearson correlation coefficient, or Jaccard index, is a widely used metric for identifying quality issues in transcriptomics, proteomics, and metabolomics data^57, 62, 82^. However, our study demonstrated the limitations of such single-sample based metrics. In particular, a high Pearson correlation coefficient between technical replicates from one single sample did not assure high quality in detecting the intrinsic biological differences between different groups of samples (**Extended Data Fig. 5**). Under such scenarios, unfortunately, the inter-sample differences between different groups of samples (*i.e*., “signal”) and the intra-sample differences of technical replicates of the same sample (*i.e*., “noise”) are at the same level, indicating that the measurement system does not have any differentiating ability. The Quartet multi-sample based reference materials suites and the SNR metric offer indispensable advantages for reliability assessment for each type of omics profiling.

Our results urge a paradigm shift from “absolute” to “ratio"-based profiling by incorporating universal reference materials in designing and executing a multiomics study. A striking finding of our study is that the multiomics profiling data at the “absolute” level, such as FPKM in transcriptomics, FOT (fraction of total) in MS-based proteomics, and relative peak areas in metabolomics from a single sample, is inherently irreproducible across platforms, labs, or batches, leading to the notorious “batch effects”. Such batch effects, usually confounded with study factors of interests, hinder the discovery of reliable biomarkers either by mistaking batch differences as biological signals or by attenuating biological signals with the incorrect use of “batch-effect correction” methods (see details in an accompanying paper^70^). The presence of batch effects makes the horizontal integration of diverse datasets from the same omics type impossible, as can be seen from the lack of capability of correctly clustering the Quartet samples (**Fig. 3a** top). Convincingly, by converting absolute profiling data of study samples to ratio scales relative to those of the same reference material (such as D6), the resulting ratio-based profiling data (such as D5/D6) were comparable across different protocols, instruments, labs, or batches (**Fig. 3a** bottom), and therefore were defined as the quantitative reference datasets (**Extended Data Fig. 4**).

The large differences in reproducibility between absolute- and ratio-based profiling data can be explained, at least partially, by the fundamental principles and assumptions behind data representation of omics measurements. The concentration or abundance of an analyte (*C*) in a sample is important to biomedical research and what a measurement technology intends to provide. In quantitative omics profiling, the “absolute” instrument readout or intensity (*I*, *e.g*., FPKM, FOT, or peak area, whether per sample scaling or normalization is applied or not) is usually used as a surrogate of *C* by assuming that there is a linear and fixed relationship (*f*, or sensitivity) between *I* and *C* under any experimental conditions^83^, *I*=*f*(*C*). In reality, however, the relationship *f* can vary due to the differences in platform details, reagent lots, lab conditions, or operator biases, among other experimental factors, making *I* inherently irreproducible between batches. On the contrary, when a common reference sample (r) is analyzed in parallel with study samples in the same experiment (batch) as a control, the resulting ratio of *I*^s^ / *I*^r^ from each batch will remain reproducible and accurately reflect the ratio of *C*^s^ / *C*^r^. It is because the intensity *I* for the reference and study samples can be represented as *I*^r^_1_=*f*_1_(*C*^r^) and *I*^s^_1_=*f*_1_(*C*^s^) for batch 1 and *I*^r^_2_=*f*_2_(*C*^r^) and *I*^s^_2_=*f*_2_(*C*^s^) for batch 2, respectively. Note that *f* remains fixed or comparable for both the reference and study samples being analyzed under the same experiment (batch). Thus, when we divide the intensity *I* of the study sample by that of the reference sample in the same batch, the resulting ratio, *I*^s^_1_ / *I*^r^_1_ for batch 1 and *I*^s^_2_ / *I*^r^_2_ for batch 2, will remain the same and equal to *C*^s^ / *C*^r^, a constant of biological significance. In fact, the lack of reproducibility of absolute gene-expression data in microarray^55, 84^, RNAseq^54^, or miRNAseq^82^ across batches or platforms have been widely documented, as is the increased reproducibility at the ratio scale^54, 55, 83^. Ironically, mainstream practices still represent omics profiling data in absolute scale, presumably due to the lack of readily accessible reference materials as controls, leading to numerous challenges in integrating diverse datasets generated under various experimental conditions. It is gratifying to note that the Olink proteomics platform reports profiling data in ratio scales relative to its control samples (www.olink.com/).

Multiomics profiling is an integrated process, and performance validation should be conducted in the entire sample-to-result process. We observed that each component of the data generation and data integration procedures can affect the final results of multiomics profiling. For each type of omics data generation, a full performance validation and proficiency testing should be conducted to assess whether the measurement can identify the inherent biological differences between various sample groups, a fundamental goal of multiomics profiling. Previous studies mainly focused on performance validation of new technologies^52, 60^, but our study revealed that horizontal and vertical data integration across technologies should also be assessed using ground-truth based objective QC metrics. The multiomics design of the Quartet Project allowed us to demonstrate the advantages of multiomics profiling over any single omics type and to objectively evaluate the pros and cons of various data integration methods in terms of clustering samples according to built-in between-group differences and identifying reliable features with cross-omics relationships obeying the central dogma rule. The Quartet Project established a novel framework for developing multiomics reference materials, reference datasets, and QC methods for multiomics studies along with the best practice guidelines for QC and data integration of multiomics profiling (**Table 2**).

Several limitations and caveats of our study should be pointed out. First, the number of analytes (*e.g*., mRNAs or proteins) expressed in the Quartet reference materials is limited. Each Quartet reference material was derived from a single B-lymphoblastoid cell line, thus genes or proteins not expressed in the B-lymphoblastoid cell line are not expected to be detectable in the Quartet reference materials. This is not a serious problem when the purpose is to use the Quartet reference materials for proficiency testing or internal optimization of technology platforms or training of lab technicians. However, this could become a limitation when the Quartet reference materials are to be used as controls and profiled along with study samples for reporting ratio-based profiling data, because the denominator for the non-detectable features would become zeros. In this case, a fudge factor or flooring value can be added to make the division possible. Secondly, the number of analytes with well-defined reference values of differential expression (ratio) between sample pairs is also limited, because only ratio values large enough are reproducibly detectable. Thirdly, although the stability of DNA is commonly accepted and the stability of MAQC reference RNA samples has demonstrated for at least 17 years (unpublished data), the long-term stability of the Quartet protein and metabolite reference materials needs to be monitored in terms of both the stability of individual analytes and the stability of the ratio-based reference values. Finally, as is true for any reference materials, the replication of the Quartet multiomics reference materials will require the recalibration of the reference datasets, and batch-to-batch differences in the production and characterization of the reference materials need to be carefully recorded and reported, such as potential genetic drifts and variability in quantitative omics features at the RNA, protein, and metabolite levels due to cell culturing.

In summary, the Chinese Quartet Project provides the international community with rich multiomics resources, which can serve as a reference for the research community to evaluate new technologies, labs, assays, products, lab operators, and computational algorithms. Large-scale multiomics studies usually involve complex multi-center and long-term measurements. To ensure the reliability of scientific research results, we highly recommend the use of unified Quartet reference materials or equivalents during generation, analysis, and integration of heterogeneous datasets. In particular, the ratio-based paradigm-shift approach using common references as side-by-side controls, when widely adopted, can fundamentally advance the integration of diverse multiomics datasets from research and the clinic by making them inherently reproducible and batch-effect resistant, hence increasing the chance of discovering reliable biomarkers for realizing precision medicine.

## Methods

### Human subjects

This study was approved by the Institutional Review Board (IRB) of the School of Life Sciences, Fudan University (BE2050). It was conducted under the principles of the Declaration of Helsinki. Four healthy volunteers from a family Quartet, as part of the Taizhou Longitudinal Study in Taizhou, Jiangsu, China were enrolled and their peripheral blood was collected to establish the human immortalized B-lymphoblastoid cell lines. All four donors signed informed consent forms.

### Establishment of the Quartet B-lymphoblastoid cell lines

We adopted the widely used protocol of using Epstein-Barr virus (EBV) to establish immortalized lymphoblastoid cell lines (LCLs) (Wheeler and Dolan, 2012). Peripheral blood mononuclear cells (PBMCs) were isolated using a lymphocyte separation solution (Ficoll). Naïve B cells were sorted by EasySep Human naïve B Cell Enrichment Kit (STEMCELL, Catalog#19254), and infected by Epstein-Barr virus (EBV) by centrifugation at 2000 rpm for 1 hour. After incubation, the successfully infected and immortalized cells were propagated in culture medium.

### Cell culture

The Quartet LCLs were cultured in RPMI 1640 with 2 mM L-glutamine, 10% heat-inactivated FBS (fetal bovine serum), and 1% PS (penicillin/streptomycin) at 37 °C with 5% CO_2_. The cells were passaged every 72 hours at a 1:4 split ratio.

### Preparation of the first batch of DNA reference materials

To obtain the first batch of DNA reference materials (Lot NO 20160806), 2 × 10^9^ cells were harvested simultaneously for each cell line. Specifically, the cells grew in suspension and were centrifuged at 300 g for 5 mins to obtain cell pellets. The cell pellets were then washed twice with cold PBS.

The DNA reference materials were purified using the DNA with Blood & Cell Culture DNA Maxi Kit (Qiagen, Germany) according to the manufacturer’s instructions, divided into 1000 aliquots for each of the Quartet members, and then labeled as Quartet_DNA_D5_20160806, Quartet_DNA_D6_20160806, Quartet_DNA_F7_20160806, and Quartet_DNA_M8_20160806. A single vial contains approximate 10 μg of genomic DNA (220 ng/μL, 50 μL) in TE buffer (10 mM TRIS, pH 8.0, 1 mM EDTA, pH 8.0).

The DNA integrity and long-term stability were evaluated by Agilent 2200 TapeStation system (Agilent Technologies, USA). Concentrations were determined by NanoDrop ND-2000 spectrophotometer (Thermo Fisher Scientific, USA).

### Preparation of multiomics reference materials

To obtain the second batch of multiomics reference materials (Lot NO 20171028), 1 × 10^10^ cells were harvested for each cell line.

2 × 10^9^ cells were used for preparing the second batch of DNA reference materials (Lot NO 20171028) with the same method mentioned above for the first batch of DNA. The second batch of DNA reference materials were stocked in 1000 vials (220 ng/μL, 50 μL), and labeled with Quartet_DNA_D5_20171028, Quartet_DNA_D6_20171028, Quartet_DNA_F7_20171028, and Quartet_DNA_M8_20171028. The DNA QC and monitoring of stability were conducted using the same methods mentioned above.

2 × 10^9^ cells pretreated with TRIzol reagent were used for preparing RNA reference materials using RNeasy Maxi kit (Qiagen, Germany) according to the manufacturer’s instructions. The extracted RNA was divided into 1000 aliquots for the quartet members and labeled as Quartet_RNA_D5_20171028, Quartet_RNA_D6_20171028, Quartet_RNA_F7_20171028, and Quartet_RNA_M8_20171028. A single vial contains approximately 5 μg of RNA in water (520 ng/μL, 10 μL). The RNA integrity and long-term stability were assessed by a 2100 Bioanalyzer using RNA 6000 Nano chips (Agilent Technologies, USA) and a Qsep 100 system (BiOptic Inc., Taiwan, China). Concentrations were determined by NanoDrop ND-2000 spectrophotometer (Thermo Fisher Scientific, USA).

2 × 10^9^ cell pellets were used for preparing protein reference materials. Two batches of peptides were prepared separately at Fudan University (on Nov 6^th^, 2017) and Novogene (on Jun 16^th^, 2020), China. Briefly, cells were lysed in 8 M urea lysis buffer supplemented with protease inhibitors. The extracted proteins were then digested using trypsin overnight at 37 °C. The peptides were divided into 1000 aliquots and dried under vacuum for each batch of peptide reference materials. Four chemically synthesized peptides with C13 and N15 labeled in valine at fixed weight ratios were spiked in the second batch of the reference protein materials (Lot NO 20200616) as external controls. The spiked peptides are YILAGVENSK (1:1,000), ADVTPADFSEWSK (1:3,000), DGLDAASYYAPVR (1:9,000), and DSPSAPVNVTVR (1:27,000).

1 × 10^9^ cell pellets were used for preparing metabolite reference materials. Briefly, cells were extracted using methanol:water = 6:1 solution. Ten xenobiotics were spiked in at known amount in each vial as external controls. They are Indoleacetic acid (25 pmol), Taurocholic acid (1 pmol), Glycocholic acid (5 pmol), Cholic acid (25 pmol), Tauroursodeoxycholic acid (2.5 pmol), Taurodeoxycholic acid (7.5 pmol), Glycoursodeoxycholic acid (1 pmol), Glycodeoxycholic acid (0.5 pmol), Ursodeoxycholic acid (25 pmol), Deoxycholic acid (50 pmol), and Sulfadimethoxine (5 pmol). The cell exacts were divided in to 1000 vials and then dried under vacuum (Labconco, USA) to obtain the cell extracts as metabolomics reference materials. Therefore, each vial contains dried cell extracts from approximately 10^6^ cells. The stability was monitored by a P300 targeted metabolomics using a UPLC-MS/MS system in Human Metabolomics Institute, Inc. (Shenzhen, China).

### Whole-genome short-read sequencing data

#### Data generation

In order to evaluate the intra-lab performance of whole-genome short-read sequencing, three replicates for each of the Quartet DNA samples were sequenced in a fixed order (D5_1, D6_1, F7_1, M8_1, D5_2, D6_2, F7_2, M8_2, D5_3, D6_3, F7_3, and M8_3). A total of 108 libraries from six labs with either PCR or PCR-free protocol were used in this study. The libraries were sequenced on short-read platforms, including Illumina HiSeq XTen, Illumina NovaSeq, MGI MGISEQ-2000, and MGI DNBSEQ-T7. In paired-end mode, the sequencing depth was at least 30×. More information was detailed in the DNA accompanying paper (Ren et al., 2022).

#### Short-read sequencing read mapping and small variants calling

The read sequences were mapped to GRCh38 (https://gdc.cancer.gov/about-data/gdc-data-processing/gdc-reference-files). Sentieon v2018.08.01 (https://www.sentieon.com/) was used to analyze raw fastq files to GVCF files. The workflow includes reads mapping by BWA-MEM, duplicates removing, indel realignments, base quality score recalibration (BQSR), and variants calling by HaplotyperCaller in GVCF mode. We used default settings for all the processes.

#### Feature encoding for small variants

To perform the calculation of SNR values and vertical integration with other quantitative omics, we used the encoding scheme for the genotypes of single nucleotide variants (SNVs). For each genomic locus, we counted all alleles occurring in a total of 108 samples from nine batches and then encoded them. Heterozygotes that were consistent with the reference genome were encoded as 0, and the others were encoded as 1. Furthermore, we used chromosome 1 to represent the whole genome for the analysis.

### Whole-genome long-read sequencing data

#### Data generation

We evaluated the performance of structural variant detection using different data analysis pipelines, without considering the technical variation from library preparation. A total of 12 libraries from three long-read sequencing platforms were generated for the Quartet DNA reference materials (one replicate for each sample). The long-read sequencing platforms used are Oxford NanoPore PromethION (~100x), PacBio Sequel (~100x), and PacBio Sequel II (~30x).

#### Long-read sequencing read mapping and structural variants calling

Reads were mapped to GRCh38 (GCA 000001405.15) from UCSC Genome Brower (http://hgdownload.soe.ucsc.edu/goldenPath/hg38/chromosomes/). Three mappers (NGMLR, minimap2 and pbmm2) and five callers (cuteSV, NanoSV, Sniffles, pbsv and SVIM) were used to call SVs.

### DNA methylation data

#### Data generation

In order to evaluate the intra-lab performance of DNA methylation, three replicates for each of the Quartet sample groups were assayed in a fixed order (D5_1, D6_1, F7_1, M8_1, D5_2, D6_2, F7_2, M8_2, D5_3, D6_3, F7_3, and M8_3). A total of 72 libraries from three labs with two different protocols and Illumina EPIC Human Methylation microarray were used in this study. More information was detailed in the Quartet Data Portal (http://chinese-quartet.org/).

#### Preprocessing of methylation data

Raw idat files were processed using R package ChAMP v2.20.1^85^ and minfi v 1.36.0^86^. The single-sample Noob (ssNoob) method^87, 88^ was used to correct for background fluorescence and dye-bias. Next, samples with a proportion of failed probes (probe detection p-value > 0.01) above 0.1 were discarded. Probes that failed in more than 10% of the remaining samples were removed. Probes with <3 beads in at least 5% of samples per probe were also removed. All non-CpG probes, SNP-related probes, multi-hit probes, probes located in chromosomes X and Y were filtered out. After preprocessing, the methylation dataset contained 735,296 probes. Finally, the corrected Meth and Unmeth signals were used to calculate M values and β values. In this process, the offset was set to 100 and the beta threshold to 0.001.

### Whole transcriptome sequencing data

#### Data generation

In order to evaluate the intra-lab performance of whole transcriptome sequencing, three replicates for each of the Quartet sample groups were sequenced in a fixed order (D5_1, D6_1, F7_1, M8_1, D5_2, D6_2, F7_2, M8_2, D5_3, D6_3, F7_3, and M8_3). A total of 252 libraries from eight labs with either poly-A selection or rRNA-removal protocol were used in this study. On average, 100 million read-pairs per replicate were sequenced on Illumina NovaSeq or MGI DNBSEQ-T7. More information was detailed in the RNA accompanying paper (Yu et al., 2022).

#### Alignment and RNA quantification

HISAT2 v2.1 was used for read alignment to the GRCh38 (version: GRCh38_snp_tran, https://genome-idx.s3.amazonaws.com/hisat/grch38_snptran.tar.gz)^89^. SAMtools v1.3.1 was used to sort and convert SAM to BAM format^90^. StringTie v1.3.4 was used for gene quantification using Ensembl reference annotation (Homo_sapiens.GRCh38.93.gtf)^91^. Ballgown v2.14.1 and prepDE.py (https://ccb.jhu.edu/software/stringtie/dl/prepDE.py) were used to produce gene expression matrix in Fragments Per Kilobase of transcript per Million mapped reads (FPKM) for downstream analysis.

### miRNA sequencing data

#### Data generation

In order to evaluate the intra-lab performance of miRNA sequencing, three replicates for each of the Quartet sample groups were sequenced in a fixed order (D5_1, D6_1, F7_1, M8_1, D5_2, D6_2, F7_2, M8_2, D5_3, D6_3, F7_3, and M8_3). A total of 72 libraries from three labs with six different protocols were used in this study. Illumina NovaSeq or HiSeq 2500 was used to generate the miRNAseq data. More information was detailed in the Quartet Data Portal (http://chinese-quartet.org/).

#### Alignment and miRNA quantification

The extra-cellular RNA processing toolkit (exceRpt) was used to pre-process miRNAseq^92^ data. The raw reads were aligned to the hg38 genome and transcriptome of exceRptDB. Counts per million mapped reads (CPM) quantifications of miRNA were extracted for the downstream analysis.

### Mass spectrometry (MS)-based proteomics data

#### Data generation

With the first batch of peptide reference materials, 312 libraries based on the LC-MS system were generated under a data-dependent acquisition mode (DDA). Samples were analyzed in a random order for each dataset, which contains three technical replicates for each of the four biological samples (D5, D6, F7, and M8). Mass spectrometers from three platforms were used: 1) Q Exactive hybrid quadrupole-Orbitrap series (Q Exactive, Q Exactive Plus, Q Exactive HF and Q Exactive HF-X), Orbitrap Fusion Tribrid series (Fusion and Fusion Lumos), Orbitrap Exploris 480 (all from Thermo Fisher Scientific, Waltham, MA, USA); 2) Triple-TOF 6600 (from SCIEX, Foster City, CA, USA), and 3) timsTOF Pro (from Bruker Daltonics, Bremen, Germany).

The second batch of peptide reference materials were all analyzed in a fixed order (D5_1, D6_1, F7_1, M8_1, D5_2, D6_2, F7_2, M8_2, D5_3, D6_3, F7_3, and M8_3) on Q Exactive, Q Exactive HF, Q Exactive HF-X, and Orbitrap Fusion Lumos, generating 36 libraries based on DDA mode and 36 libraries based on data independent acquisition (DIA) mode. All parameters were set according to the requirements from the manufacturers.

#### Peptide identification and protein quantification

MS raw files generated by the first batch of peptide reference materials were searched against the National Center for Biotechnology Information’s (NCBI) human RefSeq protein database (updated on 04-07-2013, 32,015 entries) using Firmiana 1.0 enabled with Mascot 2.3 (Matrix Science Inc.)^93^. MS raw files generated by the second batch of peptide reference materials were searched against UniProt (http://www.uniprot.org) (release-2021_04), using in-house pipelines from different labs (MaxQuant 1.5.3.17, Spectronaut 14.4, mProphet or Proteome Discoverer 2.2). Fixed modification is Carbamidomethyl (C), and Variable modifications are oxidation (M) and acetyl (Protein N-term). Proteins with at least 1 unique peptide with 1% FDR at the peptide level and Mascot ion score greater than 20 were selected for further analysis. The fraction of total (FOT) values were used for downstream analysis. FOT was defined as a protein’s iBAQ divided by the total iBAQ of all identified proteins within one sample. The FOT was multiplied by 10^5^ for the ease of presentation.

### Mass spectrometry (MS)-based metabolomics data

#### Data generation

In order to evaluate the intra-lab performance of MS-based metabolomics, three replicates for each of the Quartet sample groups were profiled in a fixed order (D5_1, D6_1, F7_1, M8_1, D5_2, D6_2, F7_2, M8_2, D5_3, D6_3, F7_3, and M8_3). The dried cell extracts were re-dissolved in mobile phase in each lab, and a total of 264 libraries were generated from five labs. The non-targeted metabolomics datasets were generated using AB SCIEX Triple TOF6600 and Thermo Scientific Q Exactive mass spectrometer systems in three different labs. The targeted metabolomics datasets were generated using Waters Xevo TQ-S, AB SCIEX QTRAP 5500, and AB SCIEX QTRAP 6500+ mass spectrometers in four labs. More information was detailed in the metabolite accompanying paper (Zhang et al., 2022).

#### Compound identification and metabolite quantification

Raw data were extracted, peak-identified and QC processed using the in-house methods in each lab. Compound identification was conducted using in-house library based on the retention time/index (RI), mass to charge ratio (m/z), and MS spectral data for each metabolite. Metabolite quantification was conducted using area-under-the-curve or the concentration calculated by calibration curve using standards of each metabolite. All the expression tables of metabolomics were log2 transformed and then normalized by Z-score transformation across all metabolites for each sample.

### Construction workflow of reference datasets

In the analysis of differentially expressed features (DEFs) and cross-omics relationships, the methylation microarray data were converted to M values, miRNA data were normalized to log2CPM, and RNA data were normalized to log2FPKM, proteomics data were normalized to log2FOT, and metabolomics data were log2 transformed based on the quantitative intensity.

Intra-batch quality control was performed to minimize the influence of technical noises in the voting process. For each sample group, features that were not detected in more than one technical replicate or that had large variability (CV > 0.15 for methylation and > 0.3 for other omics) were excluded. Afterward, we constructed the reference datasets of DEFs and cross-omics feature relationships with the consensus voting approach described below.

#### Reference datasets for DEFs

Cross-batch QC was performed following the previous intra-batch QC. Features retained in more than a certain percentage (70% for Methylation, miRNA, and RNA; 30% for protein and metabolite) of batches were kept for the subsequent differential expression analysis.

For each omics type, we analyzed the DEFs between D5 and F7 (D5/F7), D5 and M8 (D5/M8), and F7 and M8 (F7/M8) within each batch using Student’s t-test. A feature was identified as differentially expressed when satisfying the criteria of *p* < 0.05 and log2 fold change ≥ 0.5 or ≤ −0.5 for miRNA, RNA, proteins, and metabolite profiling, or *p* < 0.05 and log2 fold change ≥ 2 or ≤ −2 for methylation M values. Furthermore, we determined whether a DEM was up- or down-regulated based on the positive or negative sign of the log2 fold change.

After identifying DEFs from each batch, we kept the DEFs presented in more than 70% batches with consistent regulatory directionality (up or down). Finally, we calculated the mean log2 fold changes of all the retained intra-batch DEFs as reference values.

#### Reference datasets for cross-omics feature relationships

The reference datasets contained cross-omics feature relationships between methylation and miRNA, methylation and RNA, RNA and miRNA, RNA and protein levels, protein and metabolite. We first performed feature selection to better identify biologically meaningful correlations by annotating cross-omics features to the same genes.

For the methylation probes, we converted the features from the level of probes to genes by taking the mean value on the promoter region (TSS200 or TSS1500) to characterize the methylation level. For the other omics types, we did not perform the transformation of feature values, but simply searched for associated genes. RNA profiles were associated with gene names via Ensembl ID. Target genes associated with specific miRNA in all of miRDB^94^ (prediction scores ≥ 80), miRTarBase^95^ (support type is Functional MTI), and TargetScan^96^ would be considered as plausible. The proteomics profiles were characterized at the level of gene names. Metabolites were associated with genes on the same pathway based on the HMDB database^97^.

Afterward, we exhaustively enumerated all the batch combinations of the above five cross-omics types and conducted cross-batch QC. Associated feature pairs retained in more than a certain number of batch combinations were used for subsequent correlation analysis. This threshold is determined by the product of the respective batch and rate (70% for Methylation, miRNA, and RNA; 30% for protein and metabolite) of the two types of omics being compared.

Next, we calculated the Pearson correlation coefficients for each feature pair in each batch combination of the five cross-omics types. According to the results of Pearson correlation analysis, the cross-omics relationships were classified into positive (*R* ≥ 0.5, *p* < 0.05), negative (*R* ≤ −0.5, *p* < 0.05), and none (*p* ≥ 0.05).

Finally, we preserved the cross-omics relationships with the category that account for more than 70% as the high-confidence relations. The reference Pearson correlation coefficients were the mean value of the retained data.

### Performance metrics

#### Adjusted Rand Index (ARI)

ARI is a widely used QC metric to compare clustering results against external criteria^79^. It measures the similarity of the true labels and the clustering labels while ignoring permutations with chance normalization, which means random assignments will have an ARI score close to zero. ARI is in the range of −1 to 1, with 1 being the perfect clustering. ARI is calculated based on RI as follows:

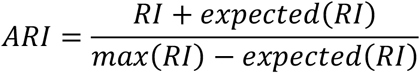

#### Root Mean Square Error (RMSE)

RMSE is the standard deviation of the residuals (prediction errors), a widely used statistic in bioinformatics and machine learning. In this study, we used RMSE to measure the consistency of DEGs detected from a dataset for a given pair of samples with those from the reference DEFs, or “RMSE of DEFs”. Reference DEFs were integrated by consensus voting the intra-batch results, and the reference difference was defined by the mean value of log2 fold change of high-confidence batches. RMSE is computed using the following equation:

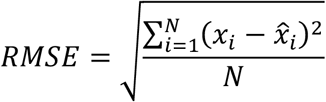

where *N* is the total number of features considered for evaluation, 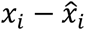 is the error, 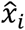 is the log2 fold change after horizontal integration, and *x*_i_ is the log2 fold change of the corresponding feature in the reference dataset.

#### Signal-to-Noise Ratio (SNR)

SNR is a parameter based on the Quartet study design for discriminating different types of reference samples. Based on principal components analysis (PCA), SNR is defined as the ratio of the average distance among different samples (like D5-1 vs. D6-1) to the average distance among technical replicates (like D5-1 vs. D5-2). SNR is calculated as follows:

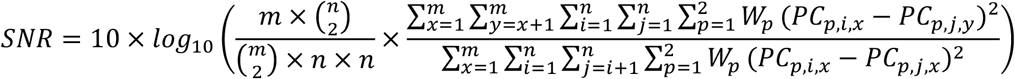

where *m* is the number of sample groups, while *n* is the number of replicates in each sample group. *W*_*p*_ represents the *p*^th^ principal component of variances. *PC*_*p*,*i*,*x*_, *PC*_*p*,*j,x*_ and *PC*_*p,j,y*_ represent the *p*^th^ component values of replicate *i* and replicate *j* in sample group *x* or sample group *y*, respectively.

### Balance of sample classes between batches

To evaluate the effect of the level of balance between the sample classes across batches on the tasks of sample classification and the identification of DEFs, we use the Jaccard index to represent the level of balance. The Jaccard index is a common statistic used for gauging the similarity and diversity of two sets.

For the Quartet multiomics datasets, the total number of samples for each of the four classes in each omic data type is the same (referred to as N). We randomly selected a natural number n from 20% to 80% of N, then we drew n samples from the sets of D5, D6, F7, and M8 and recorded the batch information from which these samples came from. Further, we calculated the Jaccard index between the batches of D5-D6, D6-F7, F7-M8, and M8-D5. Finally, the mean value of the above four Jaccard indexes represented the sample classes-batch balance.

### Within-omic (Horizontal) integration

#### Data preprocessing

First, we randomly selected four batches for each quantitative omic profiling (methylation, miRNA, RNA, protein, and metabolite). We took three D5s with one D6, three D6s with one F7, three F7s with one M8, and three M8s with one D5 from each of the four batches, to increase the difficulty of the horizontal integration task.

Next, we used different strategies for handling missing values for different omics. For methylation data, features containing missing values were removed. For other omics data, a feature will be retained when it is detected in more than 80% of the samples. For miRNAseq and RNAseq, a flooring value of 0.01 was added to each gene’s FPKM or CPM value before log2 transformation. Missing values were filled using the HM (Half of the Minimum) method.

#### Horizontal integration methods

A total of six methods were used in this study to horizontally integrate multiple batches of data, including Ratio, ComBat, Harmony, RUVg, Z-Score, and Absolute. Ratio method uses the mean value of D6 samples as the denominator to scale the expression of D5, F7, and M8 on a feature-by-feature basis. ComBat was implemented by using the ComBat function of sva v3.38.0 package^98^. Harmony was implemented by using the HarmonyMatrix function of harmony v0.1.0 package^73^. RUVg was conducted by using the RUVg function of RUVSeq v1.24.0 package^74^. Z-Score was performed by scaling each batch feature-wise before merging multiple batches to eliminate the batch effect. Absolute method refers to direct integration after normalization.

### Cross-omics (Vertical) integration

#### Two scenarios for vertical integration

(1) Integration of multi-batch quantitative omic data, related to **Fig. 4**. After horizontally integrating 16 samples from four batches of methylation, miRNA, RNA, protein and metabolite, vertical integration was performed using the five algorithms described above. (2) Integration of single-batch quantitative and genomic data, related to **Fig. 5**. The datasets of small variants were added to the integration task. Another difference is that for each omics type we only used one batch of data, which means that the vertically integrated results were not affected by problems that exist in horizontal integration, e.g, batch effects.

#### Data preprocessing

Prior to the vertical integration, we filtered out batches of very good (top 20%) or bad (bottom 20%) quality within each omics based on SNR values to reduce the impact of extreme quality datasets. In total, five batches of DNA (SNV/Indel) data, two batches of DNA methylation profiles, two batches of miRNA profiles, 11 batches of RNA profiles, 18 batches of proteomics data, and 12 batches of metabolomics data were retained.

To reduce the impact of large differences in dimensionality across multiomics on the final results, for each omics type we selected the top 1000 most variable features based on the coefficient of variation (CV). After that, the data matrices were centered and scaled to mean 0 and standard deviation 1 feature by feature. In addition, since intNMF and MCIA are methods based on the principle of non-negative matrix decomposition, features containing negative values were added with their absolute of minimum values to ensure the non-negativity.

#### Vertical integration methods

SNF^5^, iClusterBayes^75^, MOFA+^76^, MCIA^77^, and intNMF^78^ were used to integrate the multiomics data. SNFtool v2.3.1 package was used with the parameter K (number of neighbors) set to the square of the sample size after rounding, alpha (hyperparameter) to 0.05, and T (number of Iterations) to 10. iClusterPlus v1.26.0 package was used with the parameter K (number of eigen features) set to the number of sample groups minus one. MOFA2 v1.1.21 package was used with the default parameters, and PAM clustering was performed on the latent factors of the MOFA+ model to obtain the sample labels. MCIA was implemented by using omicade4 v1.30.0 package, and PAM clustering was performed on the synthetic scores to get the sample labels. IntNMF v1.2.0 package was used with the default parameters. All other parameters were set by default for the above five tools. A total of 50 iterations of data integration were performed.

### Similarity between D5 and D6

The SNF method was used to integrate data from different multiomics combinations to explore the genomic inheritance patterns of the Quartet identical twins during data integration. During integration, we randomly selected a batch from each omics with a moderate quality (SNR in the 20% to 80% range) and then calculated the inter-sample similarity matrix W using the SNFtool v2.3.1 package. Specifically, for single-omic datasets (i.e., DNA or metabolite), we treated multiple batches of moderate quality data from the same omics type as different sources, also using SNF for integration and to obtain the W matrix. As D5 and D6 each contained three technical replicates, there were nine similarity results in the W matrix. We used their mean values as the similarity between D5 and D6 obtained from one integration. To ensure the robustness of the results, a total of 50 iterations were performed for the multiomics combination.

### Statistical analysis

All statistical analyses were performed using R statistical packages (version 4.0.5) (https://www.r-project.org). Pearson’s correlation coefficients were calculated using Hmisc v4.6.0 package (https://CRAN.R-project.org/package=Hmisc). Differential expression analyses were implemented using ChAMP v2.20.1 package for methylation EPIC data^85^, and using rstatix v0.7.0 package for other omics data (https://github.com/kassambara/rstatix). PCA was conducted with the univariance scaling using the prcomp function. PAM clustering was implemented using cluster v2.1.3 package (https://CRAN.R-project.org/package=cluster). Data visualization was implemented using R packages ggplot2 v3.3.6 (https://ggplot2.tidyverse.org/), ggsci v2.9 (https://github.com/nanxstats/ggsci), ggpubr v0.4.0 (https://github.com/kassambara/ggpubr/), ComplexHeatmap v2.6.2^99^, and networkD3 v0.4 (https://christophergandrud.github.io/networkD3/).

## Materials availability

The Quartet multiomics reference materials generated in this study can be accessed from the Quartet Data Portal (https://chinese-quartet.org/) under the Administrative Regulations of the People’s Republic of China on Human Genetic Resources.

## Data and code availability

All the raw data, processed data, and reference datasets can be accessed from the Quartet Data Portal (https://chinese-quartet.org/) under the Administrative Regulations of the People’s Republic of China on Human Genetic Resources. They can also be accessed from the Genome Sequence Archive (GSA) of the National Genomics Data Center of China with BioProject ID of PRJCA007703. The source codes for the data analyses are available at https://github.com/chinese-quartet/.

## Disclaimer

This manuscript reflects the views of the authors and does not necessarily reflect those of the U.S. Food and Drug Administration. Any mention of commercial products is for clarification only and is not intended as approval, endorsement, or recommendation.

## Acknowledgements

This study was supported in part by National Key R&D Project of China (2018YFE0201603, 2018YFE0201600, and 2017YFF0204605), the National Natural Science Foundation of China (31720103909 and 32170657), Shanghai Municipal Science and Technology Major Project (2017SHZDZX01), State Key Laboratory of Genetic Engineering (SKLGE-2117), the 111 Project (B13016), and the basic research funding in key fields supported by the National Institute of Metrology of China (AKY1929 and AKYZD2202). We thank Fudan University Taizhou Institute of Health Sciences for coordinating the Taizhou Longitudinal Study and for recruiting volunteers used in the Chinese Quartet Project. Some figures in this paper were created with BioRender.com.

## Author contributions

Y.Z., L.S., X.F., J.M.L., and C.D. conceived and oversaw the study. Y.Z. and X.L. led the efforts on the immortalization of the B-lymphoblastoid cell lines. Y.Z., W.H., D.B., Z.C., B.Y.L., R.L., S.S., H.W., F.Z., X.W., P.Z., and S.Z. cultured the cell lines, isolated and purified the multiomics reference materials. Y.Z., L.S., L.D., and R.Z. coordinated (epi-)genomic data generation and analysis. Y.Z., W.H., L.Z., H.J., L.L., J.L., R.L., J.C., H.L., Z.K., D.W., F.D., H.J., S.S., H.W., R.Z., and S.Z. contributed to multiomics data generation. C.D. S.T., and Y.Z. coordinated proteomics data generation and analysis. Y.Z. coordinated metabolomics data generation and analysis. Y.Z., Y.L., J.Y., L.D., R.Z., S.T., Y.Y., L.R., W.H., Y.M., Z.C., Q.C., Q.W.C., X.C., Y.G., B.L., B.Y.L., J.L., T.Q., E.S., J.S., N.Z., P.Z., R.Z., S.Z., A.S., J.W., J.W., J.X., H.H., W.X., L.J., C.D., J.M.L., X.F., W.T. and L.S. performed data analysis and/or interpretation. J.Y. managed the datasets. Y.Z., Y.L. and L.S. wrote and revised the manuscript. All authors reviewed and approved the manuscript.

## Additional information

None.

## Extended data

**Extended data Fig. 1.**
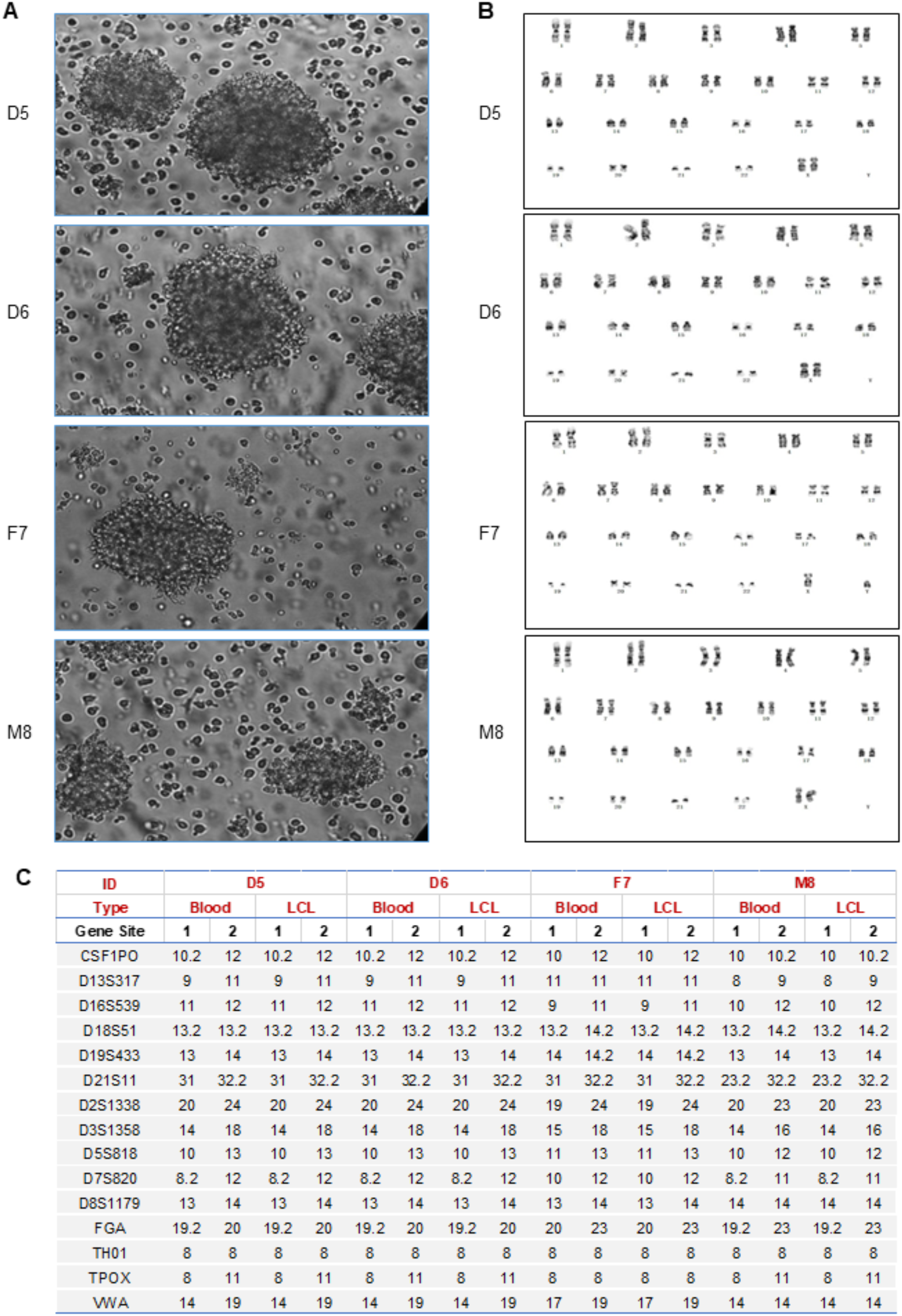
Characterization of the Quartet B-lymphoblastoid cell lines (LCLs). **a**, Quartet LCLs were cultured in suspension with typical cell clusters under phase-contrast microscopy (X400). **b**, Normal karyotypes of the LCLs were shown. **c**, 15 STR loci were used for identification of Quartet monozygotic twins’ family. Importantly, there were no differences between results from DNAs isolated from LCLs and primary blood.

**Extended data Fig. 2.**
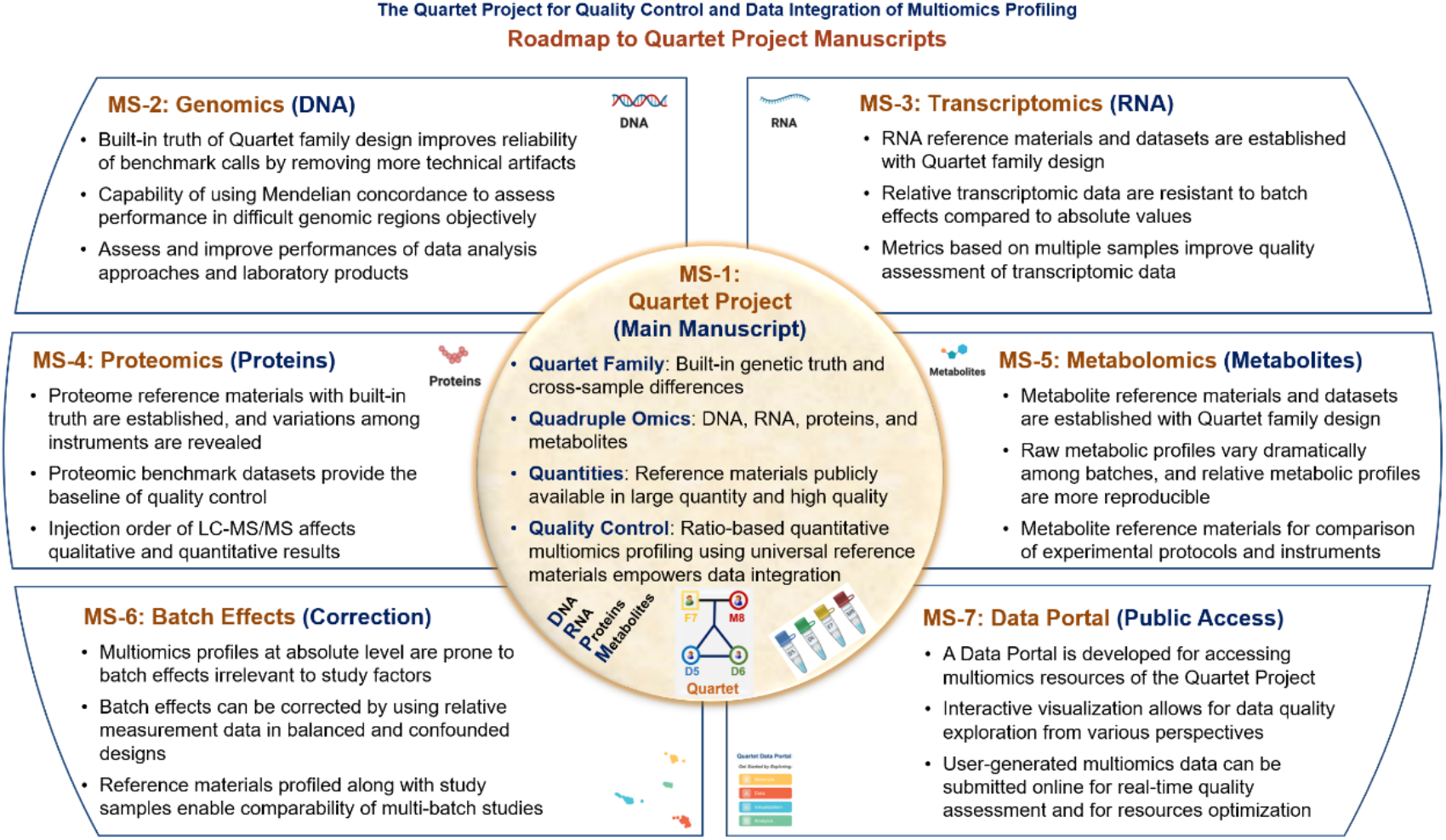
Roadmap to the Quartet Project manuscripts. **MS-1**: Quartet project overview and main findings; **MS-2/3/4/5**: Genomics/Transcriptomics/Proteomics/Metabolomics reference materials and reference datasets; **MS-6**: Batch effects and correction; **MS-7**: Data portal for public access of Quartet Project resources.

**Extended data Fig. 3.**
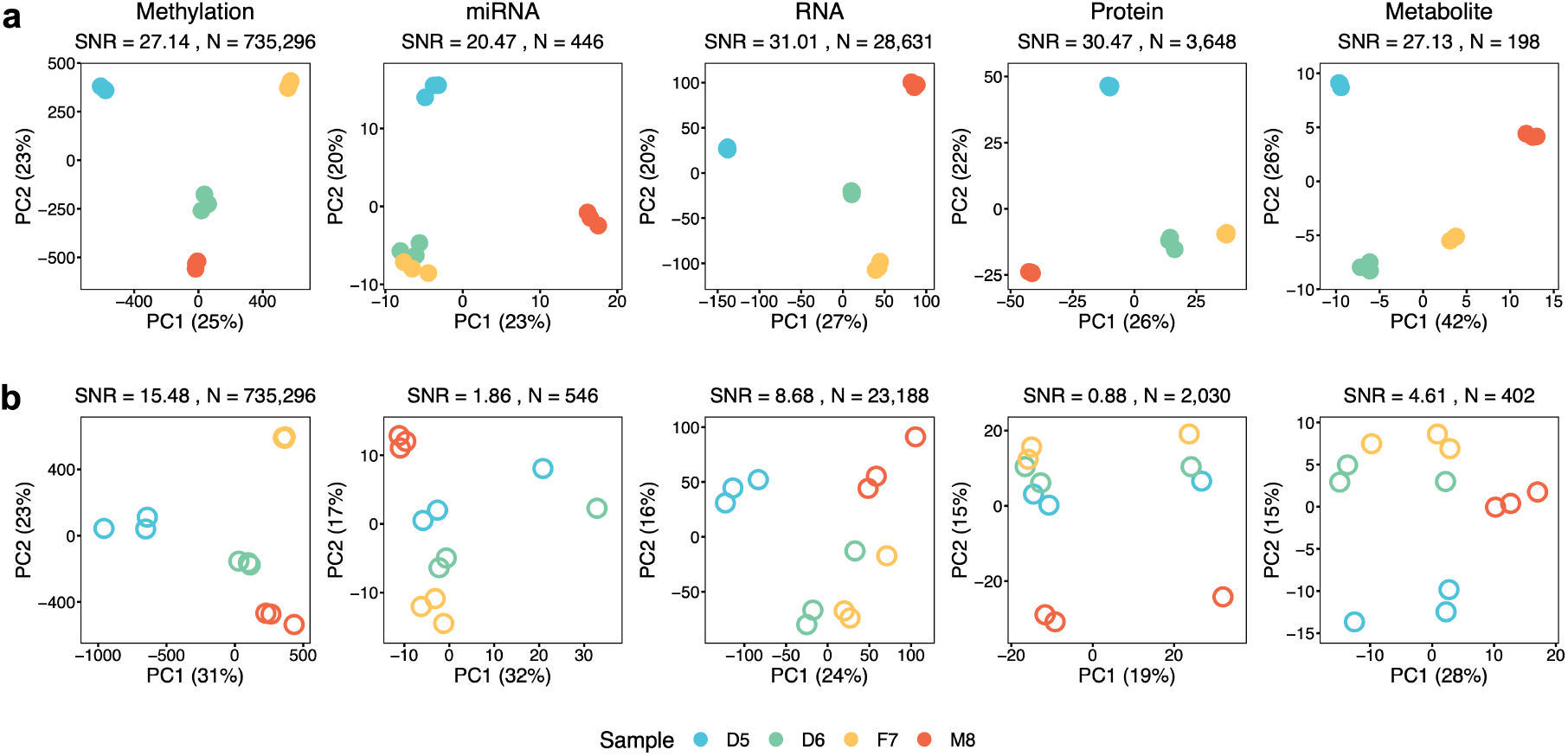
Quartet multi-sample based intra-batch Signal-to-Noise Ratio (SNR) for performance evaluation of each omics profiling. Intra-batch performance evaluation using SNR. Two batches of typically good (**a**) and bad (**b**) quality datasets of methylomics, transcriptomics, proteomics, and metabolomics were visualized by PCA plots. N is the number of features of the matrix used to calculate the SNR for the batch.

**Extended data Fig. 4.**
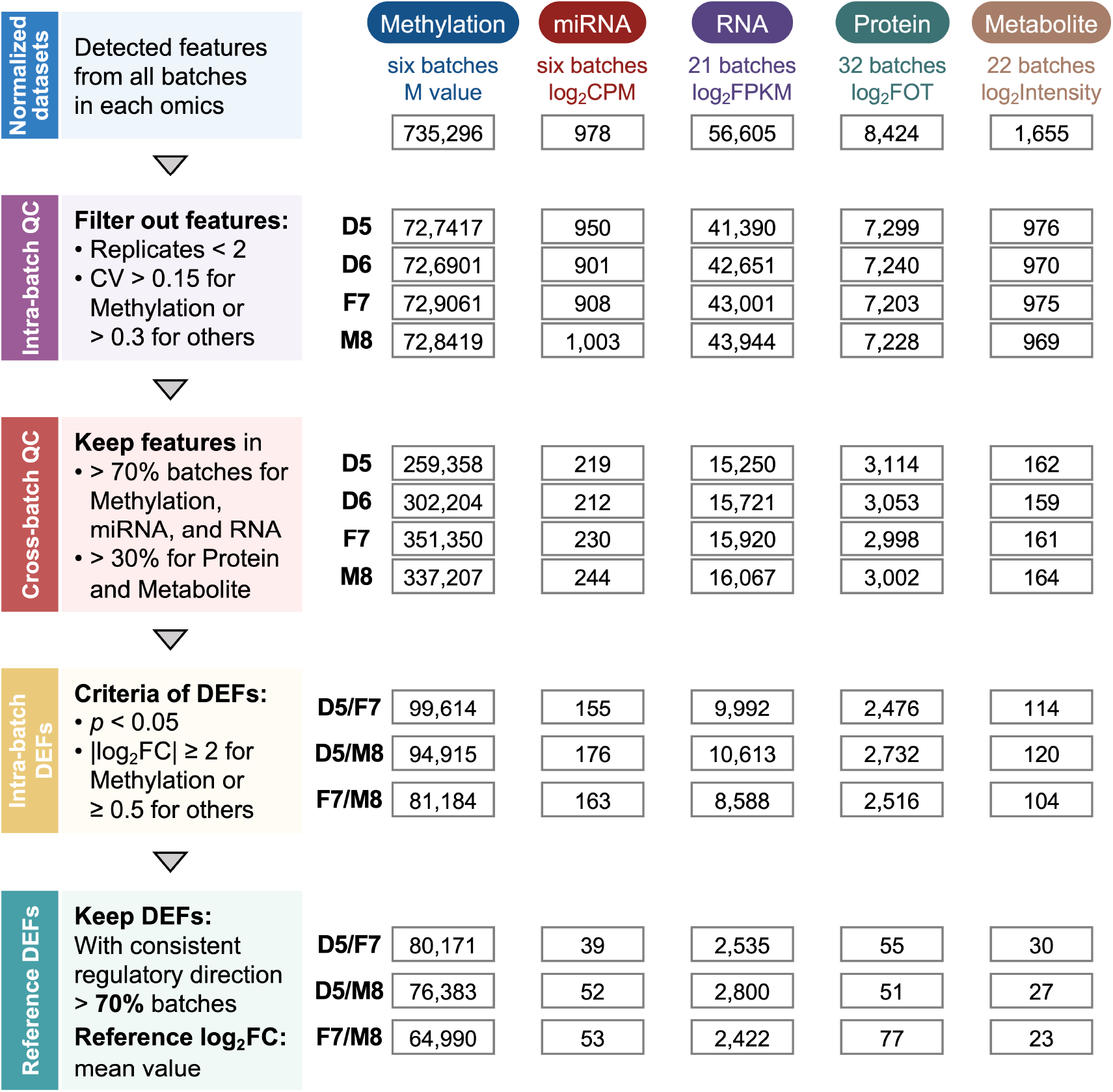
Workflow for the construction of reference datasets of differentially expressed features. Reference datasets were constructed according to the following steps: (1) Identifying detectable multiomics features and per-sample normalization. (2) Intra-batch quality control. Features that were not detectable or had low technical reproducibility were filtered out. (3) Cross-batch quality control. Features that were able to be detected in a sufficient number of batches were retained. (4) Calculating intra-batch differentially expressed features (DEFs) using t-test analysis. DEFs were classified as up- or down-regulated based on the positive or negative sign of the log2 fold change. (5) Voting based on the regulatory directionality (up or down) to screen the high confidence DEFs.

**Extended data Fig. 5.**
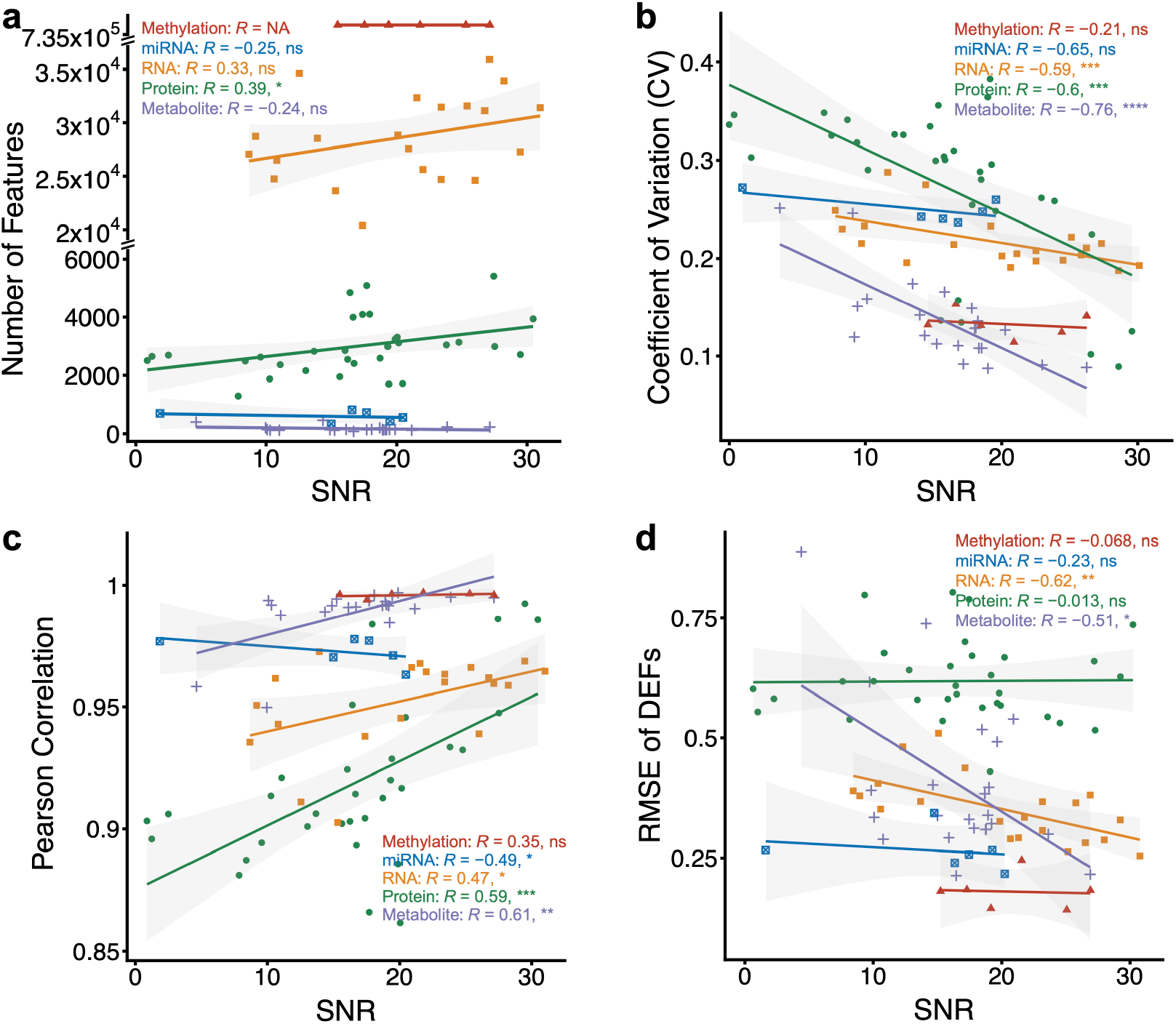
Scatter plots between SNR and the number of features, CV, Pearson correlation, and RMSE. Data points represent one batch and solid lines indicate fitted curves obtained from linear regression. Red: Methylation; Blue: miRNA; Yellow: RNA; Green: Protein; Purple: Metabolite. The annotated correlations were Pearson correlation coefficients. ns, *p* ≥ 0.05 refers to not significant, * *p*<0.05, ** *p*<0.01, *** *p*<0.001, **** *p*<0.0001. **a**, Scatter plots between SNR and number of features. **b**, Scatter plots between SNR and Coefficient of Variation (CV). The CV for each batch is the mean value of the CVs between technical replicates on all features for the four sample groups. **c**, Scatter plots between SNR and Pearson correlation coefficient, which is the mean of the results of the two-by-two calculations of the three technical replicates for each sample group. **d**, Scatter plots between SNR and RMSE of DEFs, which is the mean of RMSEs of all features within one batch.

**Extended data Fig. 6.**
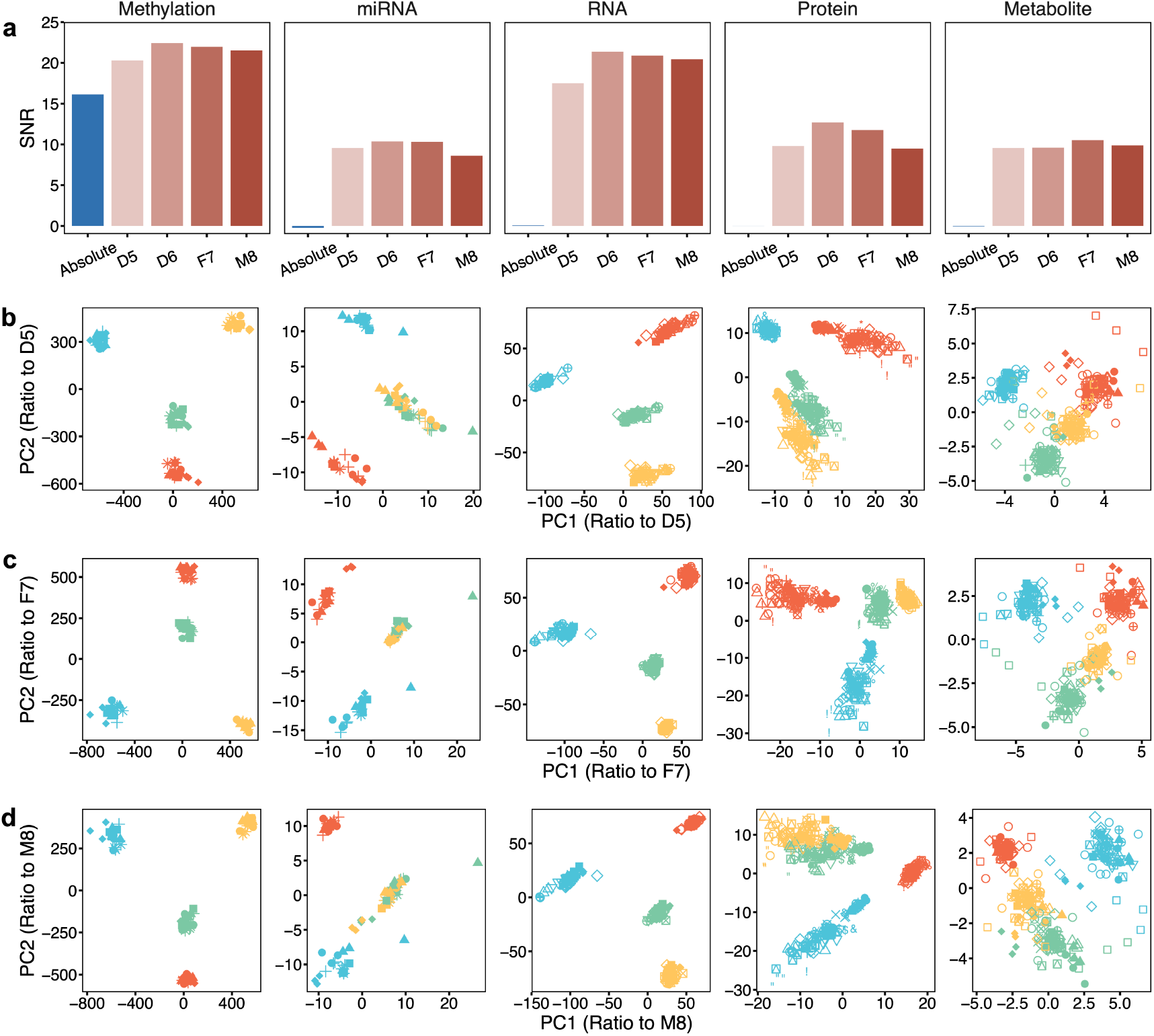
Ratio-based integration enhanced horizontal data integration and was reference-sample independent. **a**, Bar plots of Signal-to-Noise Ratio (SNR) of horizontal integration of all batches of methylation, miRNAseq, RNAseq, proteomics, and metabolomics datasets at absolute level (Blue) and ratio level (Red) with the choice of different Quartet samples as the reference sample. **b-d**, PCA plots of horizontal integration of omics datasets at ratio level by scaling to D5 (**b**), F7 (**c**), and M8 (**d**).

**Extended data Fig. 7.**
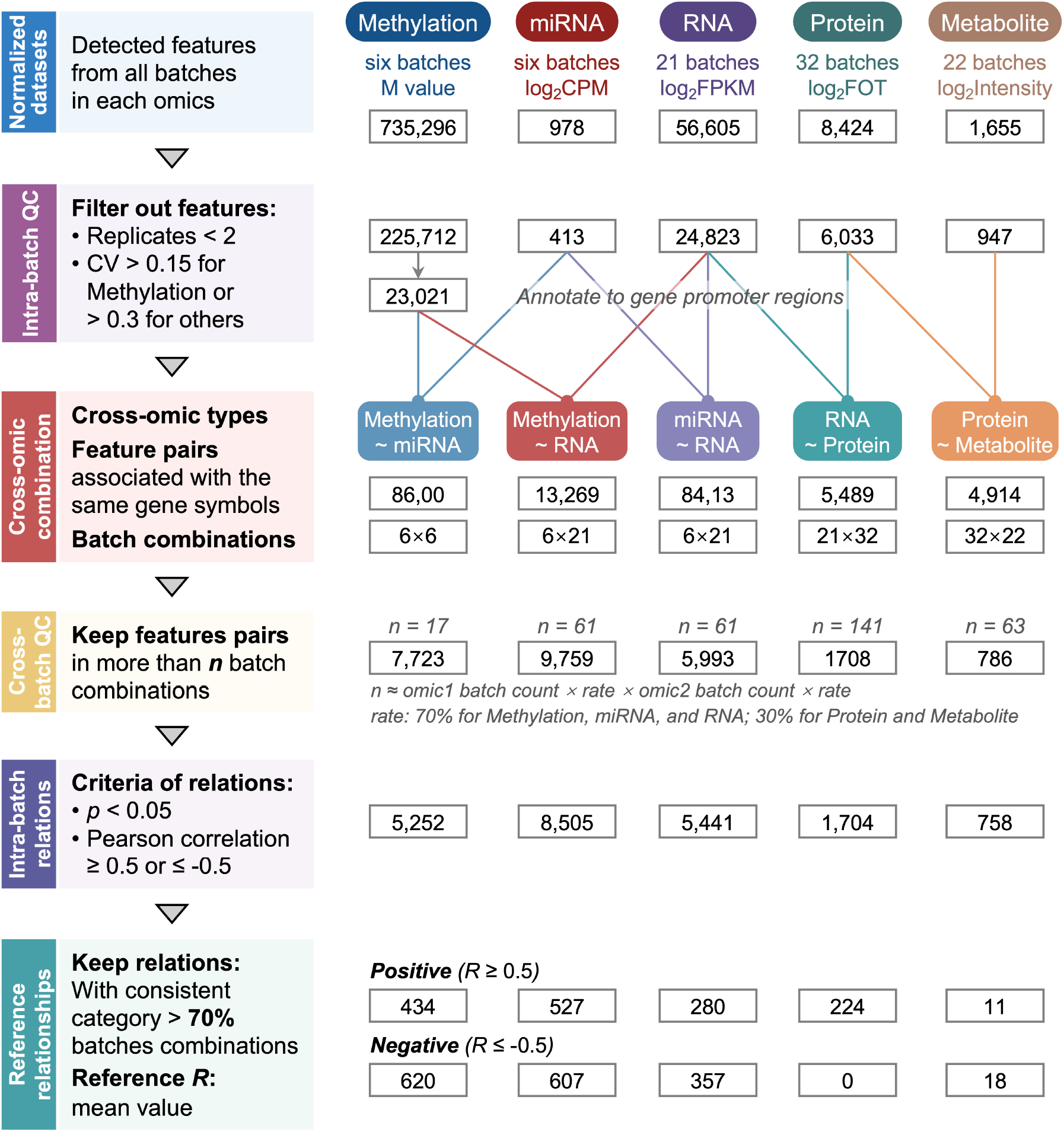
Workflow for the construction of reference datasets of cross-omics feature relationships. Reference datasets of cross-omics feature relationships were constructed according to the following steps: (1) Identifying detectable multiomics features and per-sample normalization. (2) Intra-batch quality control. Features that were not detectable or had low technical reproducibility were filtered out. (3) Identification of cross-omics feature pairs. Features associated with the same genes were retained for the five cross-omic types, i.e., methylation and miRNA, methylation and RNA, RNA and miRNA, RNA and protein, as well as protein and metabolite. (4) Cross-batch quality control. Features retained in a sufficient number of batches were used for subsequent correlation analysis. (5) Calculating Pearson correlation coefficients for each feature pair in each batch combination. Cross-omics relationships were classified into positive (*R* ≥ 0.5, *p* < 0.05), negative (*R* ≤ −0.5, *p* < 0.05), and none (*p* ≥ 0.05). (6) Voting based on the direction of the correlations (negative or positive) to screen the high-confidence cross-omics feature relationships.

**Extended data Table 1.**
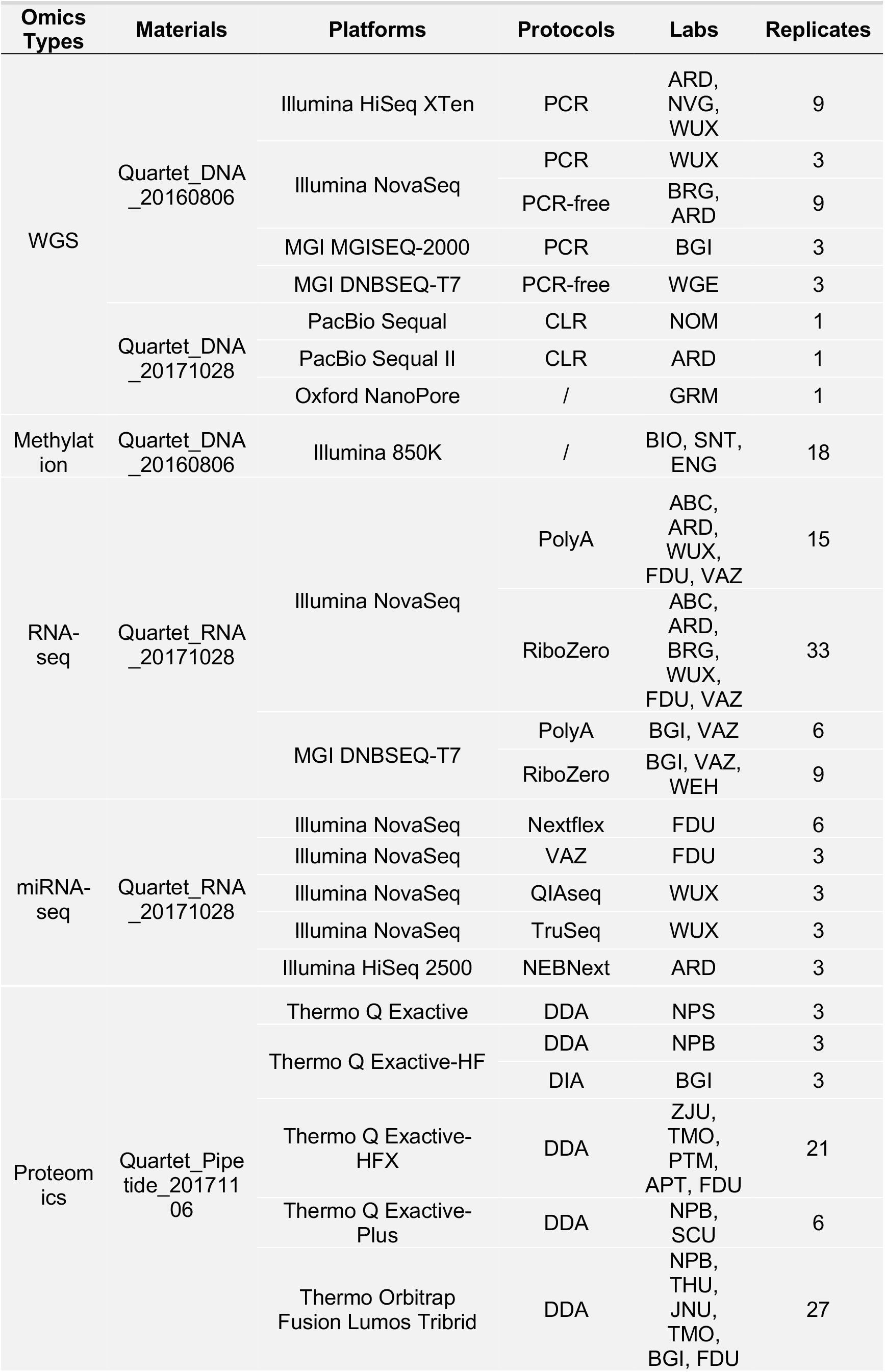
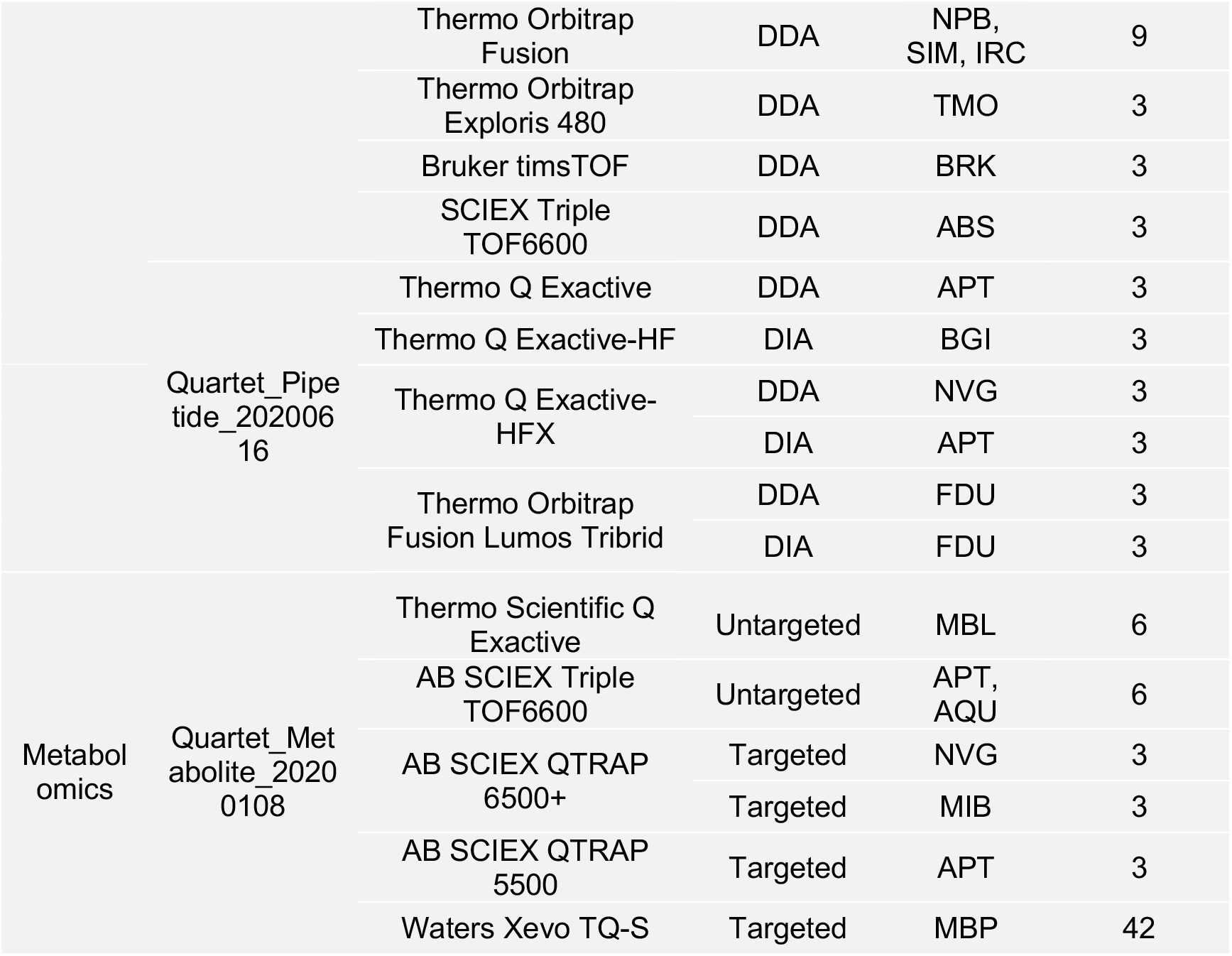
Quartet multiomics datasets generated from multiple omics types, batches, labs, and platforms.

## References

1. Hasin, Y., Seldin, M. & Lusis, A. Multi-omics approaches to disease. Genome Biol. 18, 83 (2017).

2. Karczewski, K.J. & Snyder, M.P. Integrative omics for health and disease. Nat. Rev. Genet. 19, 299–310 (2018).

3. Shilo, S., Rossman, H. & Segal, E. Axes of a revolution: challenges and promises of big data in healthcare. Nat. Med. 26, 29–38 (2020).

4. Ideker, T., Galitski, T. & Hood, L. A new approach to decoding life: systems biology. Annu. Rev. Genom. Hum. Genet. 2, 343–372 (2001).

5. Wang, B. et al. Similarity network fusion for aggregating data types on a genomic scale. Nat. Methods 11, 333–337 (2014).

6. Yan, J., Risacher, S.L., Shen, L. & Saykin, A.J. Network approaches to systems biology analysis of complex disease: integrative methods for multi-omics data. Brief. Bioinform. 19, 1370–1381 (2018).

7. Hawe, J.S., Theis, F.J. & Heinig, M. Inferring interaction networks from multi-omics data. Front. Genet. 10, 535 (2019).

8. Yurkovich, J.T., Tian, Q., Price, N.D. & Hood, L. A systems approach to clinical oncology uses deep phenotyping to deliver personalized care. Nat. Rev. Clin. Oncol. 17, 183–194 (2020).

9. Chang, K. et al. The Cancer Genome Atlas Pan-Cancer analysis project. Nat. Genet. 45, 1113–1120 (2013).

10. Bycroft, C. et al. The UK Biobank resource with deep phenotyping and genomic data. Nature 562, 203–209 (2018).

11. Campbell, P.J. et al. Pan-cancer analysis of whole genomes. Nature 578, 82–93 (2020).

12. Denny, J.C. & Collins, F.S. Precision medicine in 2030-seven ways to transform healthcare. Cell 184, 1415–1419 (2021).

13. Jin, L. Welcome to the Phenomics Journal. Phenomics 1, 1–2 (2021).

14. Veturi, Y. et al. A unified framework identifies new links between plasma lipids and diseases from electronic medical records across large-scale cohorts. Nat. Genet. 53, 972–981 (2021).

15. Tarazona, S., Arzalluz-Luque, A. & Conesa, A. Undisclosed, unmet and neglected challenges in multi-omics studies. Nat. Comput. Sci. 1, 395–402 (2021).

16. Burk, R.D. et al. Integrated genomic and molecular characterization of cervical cancer. Nature 543, 378–384 (2017).

17. Jiang, Y.Z. et al. Genomic and transcriptomic landscape of triple-negative breast cancers: subtypes and treatment strategies. Cancer Cell 35, 428–440.e425 (2019).

18. Zimmer, A. et al. The geometry of clinical labs and wellness states from deeply phenotyped humans. Nat. Commun. 12, 3578 (2021).

19. Menyhárt, O. & Győrffy, B. Multi-omics approaches in cancer research with applications in tumor subtyping, prognosis, and diagnosis. Comput. Struct. Biotechnol. J. 19, 949–960 (2021).

20. Zhou, W. et al. Longitudinal multi-omics of host–microbe dynamics in prediabetes. Nature 569, 663–671 (2019).

21. Contrepois, K. et al. Molecular choreography of acute exercise. Cell 181, 1112–1130.e1116 (2020).

22. Li, Y. et al. Using composite phenotypes to reveal hidden physiological heterogeneity in high-altitude acclimatization in a Chinese Han Longitudinal Cohort. Phenomics 1, 3–14 (2021).

23. Lehmann, B.D. et al. Multi-omics analysis identifies therapeutic vulnerabilities in triple-negative breast cancer subtypes. Nat. Commun. 12, 6276 (2021).

24. Schulte-Sasse, R., Budach, S., Hnisz, D. & Marsico, A. Integration of multiomics data with graph convolutional networks to identify new cancer genes and their associated molecular mechanisms. Nat. Mach. Intell. 3, 513–526 (2021).

25. Silverbush, D. et al. Simultaneous integration of multi-omics data improves the identification of cancer driver modules. Cell Syst. 8, 456–466.e455 (2019).

26. Price, N.D. et al. A wellness study of 108 individuals using personal, dense, dynamic data clouds. Nat. Biotechnol. 35, 747–756 (2017).

27. Tebani, A. et al. Integration of molecular profiles in a longitudinal wellness profiling cohort. Nat. Commun. 11, 4487 (2020).

28. Wilmanski, T. et al. Blood metabolome predicts gut microbiome α-diversity in humans. Nat. Biotechnol. 37, 1217–1228 (2019).

29. Dodig-Crnković, T. et al. Facets of individual-specific health signatures determined from longitudinal plasma proteome profiling. EBioMedicine 57, 102854 (2020).

30. Leiserson, M.D.M. et al. Pan-cancer network analysis identifies combinations of rare somatic mutations across pathways and protein complexes. Nat. Genet. 47, 106–114 (2015).

31. Schüssler-Fiorenza Rose, S.M. et al. A longitudinal big data approach for precision health. Nat. Med. 25, 792–804 (2019).

32. Tarazona, S. et al. Harmonization of quality metrics and power calculation in multiomic studies. Nat. Commun. 11, 3092 (2020).

33. Palsson, B. & Zengler, K. The challenges of integrating multi-omic data sets. Nat. Chem. Biol. 6, 787–789 (2010).

34. Argelaguet, R., Cuomo, A.S.E., Stegle, O. & Marioni, J.C. Computational principles and challenges in single-cell data integration. Nat. Biotechnol. 39, 1202–1215 (2021).

35. Leek, J.T. et al. Tackling the widespread and critical impact of batch effects in high-throughput data. Nat. Rev. Genet. 11, 733–739 (2010).

36. Goh, W.W.B., Wang, W. & Wong, L. Why batch effects matter in omics data, and how to avoid them. Trends Biotechnol. 35, 498–507 (2017).

37. Zhou, L., Chi-Hau Sue, A. & Bin Goh, W.W. Examining the practical limits of batch effect-correction algorithms: When should you care about batch effects? J. Genet. Genomics 46, 433–443 (2019).

38. Luecken, M.D. et al. Benchmarking atlas-level data integration in single-cell genomics. Nat. Methods 19, 41–50 (2022).

39. Tran, H.T.N. et al. A benchmark of batch-effect correction methods for single-cell RNA sequencing data. Genome Biol. 21, 12 (2020).

40. Misra, B.B., Langefeld, C.D., Olivier, M. & Cox, L.A. Integrated omics: tools, advances and future approaches. J. Mol. Endocrinol. (2018).

41. Krassowski, M., Das, V., Sahu, S.K. & Misra, B.B. State of the field in multi-omics research: From computational needs to data mining and sharing. Front. Genet. 11 (2020).

42. Cantini, L. et al. Benchmarking joint multi-omics dimensionality reduction approaches for the study of cancer. Nat. Commun. 12, 124 (2021).

43. Rappoport, N. & Shamir, R. Multi-omic and multi-view clustering algorithms: review and cancer benchmark. Nucleic Acids Res. 47, 1044 (2019).

44. Choobdar, S. et al. Assessment of network module identification across complex diseases. Nat. Methods 16, 843–852 (2019).

45. Sené, M., Gilmore, I. & Janssen, J.T. Metrology is key to reproducing results. Nature 547, 397–399 (2017).

46. Hardwick, S.A., Deveson, I.W. & Mercer, T.R. Reference standards for next-generation sequencing. Nat. Rev. Genet. 18, 473–484 (2017).

47. Salit, M. & Woodcock, J. MAQC and the era of genomic medicine. Nat. Biotechnol. 39, 1066–1067 (2021).

48. Choquette, S.J., Duewer, D.L. & Sharpless, K.E. NIST reference materials: utility and future. Annu. Rev. Anal. Chem. 13, 453–474 (2020).

49. Zook, J.M. et al. An open resource for accurately benchmarking small variant and reference calls. Nat. Biotechnol. 37, 561–566 (2019).

50. Zook, J.M. et al. A robust benchmark for detection of germline large deletions and insertions. Nat. Biotechnol. 38, 1347–1355 (2020).

51. Jones, W. et al. A verified genomic reference sample for assessing performance of cancer panels detecting small variants of low allele frequency. Genome Biol. 22, 111 (2021).

52. Deveson, I.W. et al. Evaluating the analytical validity of circulating tumor DNA sequencing assays for precision oncology. Nat. Biotechnol. (2021).

53. Fang, L.T. et al. Establishing community reference samples, data and call sets for benchmarking cancer mutation detection using whole-genome sequencing. Nat. Biotechnol. 39, 1151–1160 (2021).

54. Su, Z. et al. A comprehensive assessment of RNA-seq accuracy, reproducibility and information content by the Sequencing Quality Control Consortium. Nat. Biotechnol. 32, 903–914 (2014).

55. Shi, L. et al. The MicroArray Quality Control (MAQC) project shows inter- and intraplatform reproducibility of gene expression measurements. Nat. Biotechnol. 24, 1151–1161 (2006).

56. Friedman, D.B. et al. The ABRF Proteomics Research Group studies: educational exercises for qualitative and quantitative proteomic analyses. Proteomics 11, 1371–1381 (2011).

57. Ulmer, C.Z. et al. LipidQC: method validation tool for visual comparison to SRM 1950 using NIST interlaboratory comparison exercise lipid consensus mean estimate values. Anal. Chem. 89, 13069–13073 (2017).

58. Krusche, P. et al. Best practices for benchmarking germline small-variant calls in human genomes. Nat. Biotechnol. 37, 555–560 (2019).

59. Matthijs, G. et al. Guidelines for diagnostic next-generation sequencing. Eur. J. Hum. Genet. 24, 1515 (2016).

60. Gargis, A.S. et al. Assuring the quality of next-generation sequencing in clinical laboratory practice. Nat. Biotechnol. 30, 1033–1036 (2012).

61. Broadhurst, D. et al. Guidelines and considerations for the use of system suitability and quality control samples in mass spectrometry assays applied in untargeted clinical metabolomic studies. Metabolomics 14, 72 (2018).

62. Collins, B.C. et al. Multi-laboratory assessment of reproducibility, qualitative and quantitative performance of SWATH-mass spectrometry. Nat. Commun. 8, 291 (2017).

63. Beger, R.D. et al. Towards quality assurance and quality control in untargeted metabolomics studies. Metabolomics 15 (2019).

64. Wang, X. et al. QC metrics from CPTAC raw LC-MS/MS data interpreted through multivariate statistics. Anal. Chem. 86, 2497–2509 (2014).

65. Chen, X.D., Jiang, Y.F., Xu, P. & Jin, L. Construction and utilization of human genetic resources in large population cohorts. Yi Chuan 43, 980–987 (2021).

66. Ren, L. et al. Quartet DNA reference materials and datasets for comprehensively evaluating germline variants calling performance. bioRxiv, https://doi.org/10.1101/2022.09.28.509844 (2022).

67. Yu, Y. et al. Quartet RNA reference materials and ratio-based reference datasets for reliable transcriptomic profiling. bioRxiv, https://doi.org/10.1101/2022.09.26.507265 (2022).

68. Tian, S. et al. Quartet protein reference materials and datasets for multi-platform assessment of label-free proteomics [Unpublished manuscript]. (2022).

69. Zhang, N. et al. Quartet metabolite reference materials and datasets for inter-laboratory reliability assessment of metabolomics studies [Unpublished manuscript]. (2022).

70. Yu, Y. et al. Correcting batch effects in large-scale multiomic studies using a reference-material-based ratio method. bioRxiv, https://doi.org/10.1101/2022.10.19.507549 (2022).

71. Yang, J. et al. The Quartet Data Portal: integration of community-wide resources for multiomics quality control. bioRxiv, https://doi.org/10.1101/2022.09.26.507202 (2022).

72. Zhang, Y., Parmigiani, G. & Johnson, W.E. ComBat-seq: batch effect adjustment for RNA-seq count data. NAR Genom. Bioinform. 2 (2020).

73. Korsunsky, I. et al. Fast, sensitive and accurate integration of single-cell data with Harmony. Nat. Methods 16, 1289–1296 (2019).

74. Risso, D., Ngai, J., Speed, T.P. & Dudoit, S. Normalization of RNA-seq data using factor analysis of control genes or samples. Nat. Biotechnol. 32, 896–902 (2014).

75. Mo, Q. et al. A fully Bayesian latent variable model for integrative clustering analysis of multi-type omics data. Biostatistics 19, 71–86 (2017).

76. Argelaguet, R. et al. MOFA+: a statistical framework for comprehensive integration of multi-modal single-cell data. Genome Biol. 21, 111 (2020).

77. Meng, C., Kuster, B., Culhane, A.C. & Gholami, A.M. A multivariate approach to the integration of multi-omics datasets. BMC Bioinform. 15, 162 (2014).

78. Chalise, P. & Fridley, B.L. Integrative clustering of multi-level ‘omic data based on non-negative matrix factorization algorithm. PLoS One 12, e0176278 (2017).

79. Hubert, L. & Arabie, P. Comparing partitions. J Classif. 2, 193–218 (1985).

80. Schubert, E. & Rousseeuw, P.J. Fast and eager k-medoids clustering: O (k) runtime improvement of the PAM, CLARA, and CLARANS algorithms. Information Systems 101, 101804 (2021).

81. Baker, M. 1,500 scientists lift the lid on reproducibility. Nature 533, 452–454 (2016).

82. Giraldez, M.D. et al. Comprehensive multi-center assessment of small RNA-seq methods for quantitative miRNA profiling. Nat. Biotechnol. 36, 746–757 (2018).

83. Shi, L. et al. Microarray scanner calibration curves: characteristics and implications. BMC Bioinform. 6, S11 (2005).

84. Chen, J.J., Hsueh, H.-M., Delongchamp, R.R., Lin, C.-J. & Tsai, C.-A. Reproducibility of microarray data: a further analysis of microarray quality control (MAQC) data. BMC Bioinform. 8, 412 (2007).

## References

79. Hubert, L. & Arabie, P. Comparing partitions. J Classif. 2, 193–218 (1985).

85. Tian, Y. et al. ChAMP: updated methylation analysis pipeline for Illumina BeadChips. Bioinformatics 33, 3982–3984 (2017).

86. Aryee, M.J. et al. Minfi: a flexible and comprehensive Bioconductor package for the analysis of Infinium DNA methylation microarrays. Bioinformatics 30, 1363–1369 (2014).

87. Fortin, J.-P., Triche Jr, T.J. & Hansen, K.D. Preprocessing, normalization and integration of the Illumina HumanMethylationEPIC array with minfi. Bioinformatics 33, 558–560 (2017).

88. Triche Jr, T.J., Weisenberger, D.J., Van Den Berg, D., Laird, P.W. & Siegmund, K.D. Low-level processing of Illumina Infinium DNA methylation beadarrays. Nucleic Acids Res. 41, e90–e90 (2013).

89. Kim, D., Paggi, J.M., Park, C., Bennett, C. & Salzberg, S.L. Graph-based genome alignment and genotyping with HISAT2 and HISAT-genotype. Nat. Biotechnol. 37, 907–915 (2019).

90. Li, H. A statistical framework for SNP calling, mutation discovery, association mapping and population genetical parameter estimation from sequencing data. Bioinformatics 27, 2987–2993 (2011).

91. Pertea, M. et al. StringTie enables improved reconstruction of a transcriptome from RNA-seq reads. Nat. Biotechnol. 33, 290–295 (2015).

92. Rozowsky, J. et al. exceRpt: a comprehensive analytic platform for extracellular RNA profiling. Cell Syst. 8, 352–357. e353 (2019).

93. Feng, J. et al. Firmiana: towards a one-stop proteomic cloud platform for data processing and analysis. Nat. Biotechnol. 35, 409–412 (2017).

94. Wong, N. & Wang, X. miRDB: an online resource for microRNA target prediction and functional annotations. Nucleic Acids Res. 43, D146–D152 (2015).

95. Huang, H.-Y. et al. miRTarBase 2020: updates to the experimentally validated microRNA–target interaction database. Nucleic Acids Res. 48, D148–D154 (2020).

96. McGeary, S.E. et al. The biochemical basis of microRNA targeting efficacy. Science 366, eaav1741 (2019).

97. Wishart, D.S. et al. HMDB 5.0: the human metabolome database for 2022. Nucleic Acids Res. 50, D622–D631 (2022).

98. Leek, J.T., Johnson, W.E., Parker, H.S., Jaffe, A.E. & Storey, J.D. The sva package for removing batch effects and other unwanted variation in high-throughput experiments. Bioinformatics 28, 882–883 (2012).

99. Gu, Z., Eils, R. & Schlesner, M. Complex heatmaps reveal patterns and correlations in multidimensional genomic data. Bioinformatics 32, 2847–2849 (2016).

